# Peptidoglycan recycling contributes to outer membrane integrity and carbapenem tolerance in *Acinetobacter baumannii*

**DOI:** 10.1101/2021.11.23.469614

**Authors:** Nowrosh Islam, Misha I. Kazi, Katie N. Kang, Jacob Biboy, Joe Gray, Feroz Ahmed, Richard D. Schargel, Cara C. Boutte, Tobias Dörr, Waldemar Vollmer, Joseph M. Boll

**Author notes:** Contributed equally (Nowrosh Islam, Misha I. Kazi). Correspondence: Joseph M. Boll.

## Abstract

The Gram-negative cell envelope is an essential structure that not only protects the cell against lysis from the internal turgor, but also forms a barrier to limit entry of antibiotics. Some of our most potent bactericidal antibiotics, the β-lactams, exploit the essentiality of the cell envelope by inhibiting its biosynthesis, typically inducing lysis and rapid death. However, many Gram-negative bacteria exhibit “antibiotic tolerance”, the ability to sustain viability in the presence of β-lactams for extended time periods. Despite several studies showing that antibiotic tolerance contributes directly to treatment failure, and is a steppingstone in acquisition of true resistance, the molecular factors that promote intrinsic tolerance are not well-understood. *Acinetobacter baumannii* is a critical-threat nosocomial pathogen notorious for its ability to rapidly develop multidrug resistance. While typically reserved to combat multidrug resistant infections, carbapenem β-lactam antibiotics (i.e., meropenem) are first-line prescriptions to treat *A. baumannii* infections. Meropenem tolerance in Gram-negative pathogens is characterized by morphologically distinct populations of spheroplasts, but the impact of spheroplast formation is not fully understood. Here, we show that susceptible *A. baumannii* clinical isolates demonstrate high intrinsic tolerance to meropenem, form spheroplasts with the antibiotic and revert to normal growth after antibiotic removal. Using transcriptomics and genetics screens, we characterized novel tolerance factors and found that outer membrane integrity maintenance, drug efflux and peptidoglycan homeostasis collectively contribute to meropenem tolerance in *A. baumannii*. Furthermore, outer membrane integrity and peptidoglycan recycling are tightly linked in their contribution to meropenem tolerance in *A. baumannii*.

**Importance:** Carbapenem treatment failure associated with “superbug” infections has rapidly increased in prevalence, highlighting an urgent need to develop new therapeutic strategies. Antibiotic tolerance can directly lead to treatment failure but has also been shown to promote acquisition of true resistance within a population. While some studies have addressed mechanisms that promote tolerance, factors that underlie Gram-negative bacterial survival during carbapenem treatment are not well-understood. Here, we characterized a role for peptidoglycan recycling in outer membrane integrity maintenance and carbapenem tolerance in *A. baumannii*. These studies suggest that the pathogen limits antibiotic concentrations in the periplasm and highlights physiological processes that could be targeted to improve antimicrobial treatment.

## Introduction

The cell envelope is a dynamic barrier composed of an inner (cytoplasmic) membrane, a periplasm that includes a thin peptidoglycan (PG) layer and an outer membrane, which is a selective barrier that restricts entry of toxins and antibiotics. While the PG layer is known to protect against bursting due to the cell turgor, the outer membrane also protects against lysis when external osmotic conditions change^1^. Perturbation of the outer membrane or PG envelope layers induces lysis, but regulated responses that fortify the envelope can maintain envelope homeostasis to promote pathogen survival during stress exposure^2^.

Antibiotic treatment failure is a growing threat to public health and has primarily been associated with antibiotic resistance (i.e., the ability to grow in the presence of antibiotics). However, antibiotic tolerance, a population’s ability to survive otherwise toxic levels of transient antibiotic treatment for extended periods, likely acts as a stepping-stone to true resistance^3–5^. Antibiotic tolerance is characterized by survival of cell populations in a non-dividing state, where the minimal inhibitory concentration does not change and cells revert to normal growth when the antibiotic is removed, degraded or diluted^6–8^. Molecular factors that extend survival during treatment, increase the probability of resistance-conferring mutations or horizontal gene transfer to occur^3^.

Carbapenems are important β-lactam therapeutics because they possess potent broad-spectrum activity and are not susceptible to common resistance mechanisms^9, 10^. In fact, meropenem is a last-line carbapenem antibiotic used to treat multidrug resistant Gram-negative infections^11, 12^. While meropenem treatment is typically reserved to fight multidrug resistant bacteria, it is a first-line prescription against the highly drug resistant nosocomial pathogen, *Acinetobacter baumannii*^13, 14^. Carbapenem-resistant *A. baumannii* has become commonplace among hospital acquired infections. In 2019 the Center for Disease Control listed carbapenem-resistant *A. baumannii* as one of the most urgent threats to public health^15^, and a recent report by the World Health Organization prioritized the pathogen as critical for new antibiotic development^16^, underscoring the severity.

We reasoned that since tolerance is a prerequisite for true resistance, factors that promote carbapenem tolerance may be widespread among susceptible *A. baumannii* strains. Defining intrinsic tolerance factors in *A. baumannii* may offer fundamental insight into how resistance mechanisms rapidly spread among populations and provide new targets to combat tolerant pathogens. While our understanding of resistance mechanisms that cause antibiotic treatment failure has been well-documented, tolerance factors that precede acquisition of true resistance are limited.

Here, we show that susceptible *A. baumannii* strains, including laboratory-adapted strains and recent clinical isolates, survive for extended periods (>24 h) in high levels of meropenem, demonstrating widespread tolerance. Meropenem induces cell wall-deficient spheroplast formation in *A. baumannii*, as shown in other Gram-negative pathogens^17–19^. After removal of the antibiotic, cells rapidly revert to the canonical *A. baumannii* coccobacilli morphology and resume growth. Transcriptome sequencing analysis at timepoints leading to spheroplast formation showed differential expression of genes that coordinate a regulatory response to reduce the intracellular meropenem concentration. During meropenem treatment, outer membrane integrity and permeability contribute to fitness, which we show are also impacted by defects in the PG recycling pathway. PG recycling is also a major contributor to *A. baumannii* survival during meropenem treatment, where disruption of genes encoding periplasmic and cytoplasmic PG maintenance enzymes compromise outer membrane integrity. Lastly, we also define PBP7 (encoded by *pbpG*) and LdtK enzymatic activities, which are tolerance determinants in *A. baumannii.* Together, these studies show several pathways that coordinate in *A. baumannii* to limit meropenem-induced cell envelope damage. These findings provide new targets to direct antimicrobial therapies and prevent the spread of resistance.

## Results

### Meropenem susceptible *A. baumannii* strains are tolerant, form spheroplasts and resume normal morphology and growth upon removal of the bactericidal antibiotic

It was previously shown that *Vibrio cholerae*^17, 18^, *Pseudomonas aeruginosa*^19^ and pathogens in the Enterobacterales order^20, 21^ form viable, non-dividing spheroplasts when exposed to lethal concentration of β-lactam antibiotics over several hours. Importantly, spheroplasts revert to normal rod-shaped growth when the antibiotic concentration is sufficiently reduced^21^, demonstrating a short-term survival mechanism that directly contributes to antibiotic treatment failure.

To determine if populations of *A. baumannii* strains can tolerate meropenem treatment over time, stationary phase cultures from susceptible *A. baumannii* isolates, including recent clinical isolates, were treated with high levels (10 μg/mL; 62.5-fold MIC in ATCC 17978) of the antibiotic. Treated cultures demonstrated only slight depletion after 24 h, relative to untreated (Fig 1A; Fig S1A). In contrast, meropenem treatment of cells in logarithmic growth phase showed rapid lysis (Fig S1B). Therefore, *A. baumannii* strains in stasis, a relevant physiological state during infection when the cell is known to fortify the cell envelope and slow growth/division^22^, are highly tolerant to lethal meropenem concentrations. While these data agree with current dogma that β-lactam-dependent killing is strictly proportional with growth rate^7, 23, 24^, subsequent analysis revealed that stationary phase *A. baumannii* cells experience significant cell envelope damage upon meropenem treatment. After 12 h, stationary phase cells treated with meropenem demonstrated notable morphological changes typical of spheroplast formation relative to untreated cells (Fig 1B; Fig S2). All strains showed a measurable increase in surface area and width of treated cells relative to untreated (Fig 1C; Fig S2). A significant decrease in uptake of the fluorescent D-amino acid, NADA, was also evident (Fig 1D; Fig S2), suggesting degradation of the cell wall, as previously shown in other β-lactam tolerant Gram-negative bacteria^17, 21^. Thus, tolerance under stationary phase conditions is not just a simple function of lack of growth, but rather an active response to significant cell envelope damage.

**Figure 1:**
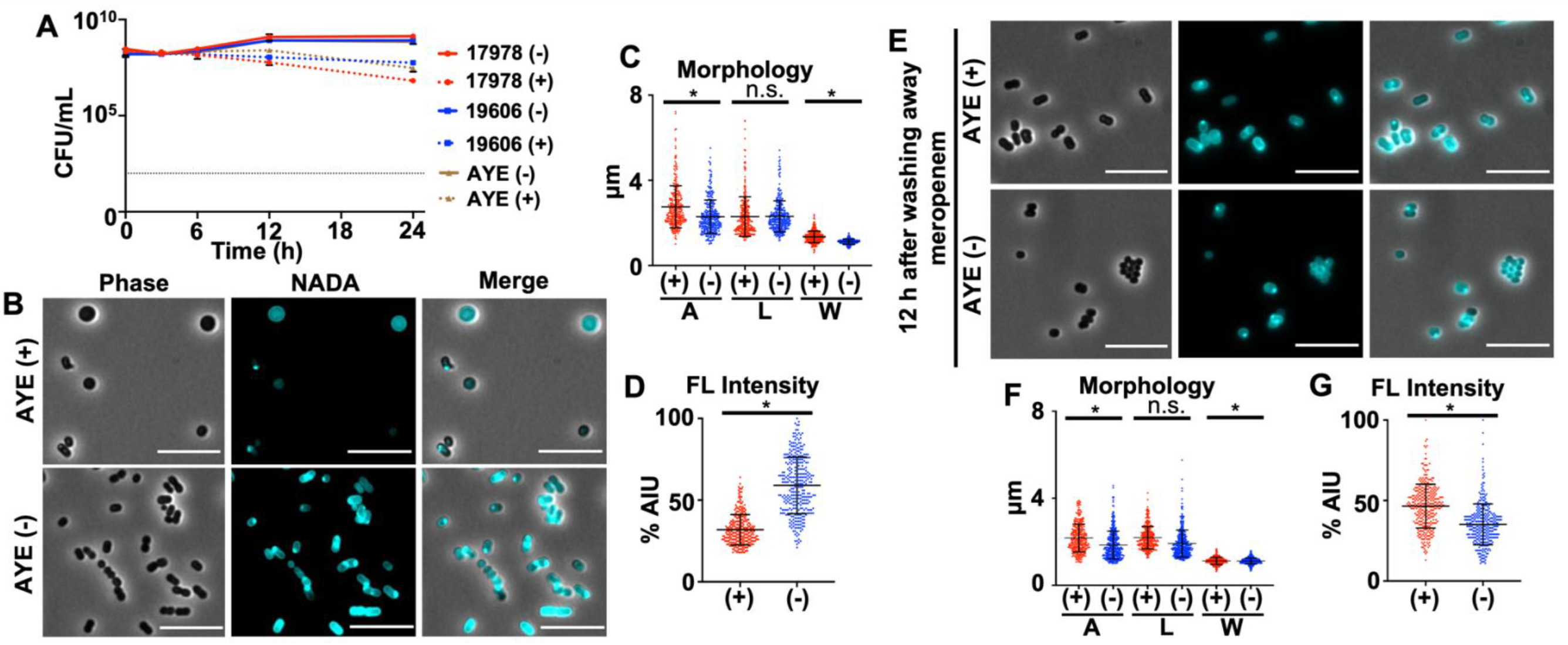
*Acinetobacter baumannii* strains are tolerant to meropenem. (A) Colony-forming units (CFUs) of *A. baumannii* strains ATCC 17978, 19606 and AYE untreated (-) or treated (+) with meropenem over 24 h. Each killing assay was independently replicated three times, and one representative dataset was reported. Dotted black line indicates level of detection. (B) Phase and fluorescence microscopy of + or -*A. baumannii* strain AYE after 12 h. Scale bar is 10 μm. (C) Area (A), length (L) and width (W) quantitation of cells in panel B (*n*= 300). (D) Fluorescent (FL) signal intensity quantitation in percent arbitrary intensity units (AIU) of treated vs. untreated cells in panel B (*n*= 300). (E) Same as panel B, but 12 hours after meropenem removal showing the characteristic *A. baumannii* coccobacilli morphology is restored. (F) Area (A), length (L) and width (W) of cells in panel E (*n*= 300). (G) FL signal intensity in percent AIU treated vs. untreated in panel E (*n*= 300). Significance was determined using an unpaired t-test (*P* <0.05) in treated vs. untreated. An asterisk indicates significant differences between treated and untreated; n.s., not significant. Error bars indicate standard deviation from the mean.

Since we showed that *A. baumannii* spheroplasts were viable after plating (Fig 1A), we also wanted to determine if the characteristic *A. baumannii* coccobacilli morphology was restored after antibiotic removal. Cells were incubated statically in fresh media without antibiotic. At 12 h post-treatment, no spheroplast were found (Fig 1E), wild type morphology was restored (Fig 1F) and the cells showed incorporation of NADA (Fig 1G). Fluorescence intensity measurements were equivalent in treated and untreated cells. Furthermore, fluorescence intensity was higher at the midcell, where the divisome regulates daughter cell formation, suggesting the recovered population had resumed division (Fig 1E). Together, these data indicate that *A. baumannii* resumes wild type morphology and growth when meropenem treatment is stopped.

### Transcriptome analysis highlights differentially regulated pathways important for *A. baumannii* tolerance

Many Gram-negative pathogens form spheroplasts within 6 h to develop meropenem tolerance^17, 21^; however, *A. baumannii* spheroplast formation is delayed. We first observe spheroplast formation only after 8 h, with large numbers within the population accumulating by 12 h (Fig 1BCD; Fig S2). To define transcriptional alterations associated with spheroplast-associated tolerance, we isolated RNA from treated and untreated cells at 0.5, 3 and 9 h. While subtle changes in gene expression were evident at 0.5 and 3 h, differential expression patterns were more obvious at 9 h in treated cultures relative to untreated (Fig 2). Genes associated with efflux were increasingly upregulated with each timepoint (Fig 2A), suggesting the cell quickly and continually responds to meropenem treatment by actively expelling the toxic compound. Upregulated efflux genes included *adeB*, *adeIJK* and *macABtolC*, which have all been implicated in antibiotic efflux^25–27^; specifically, β-lactam efflux has been reported associated with the AdeIJK RND-type pump^28, 29^.

**Figure 2:**
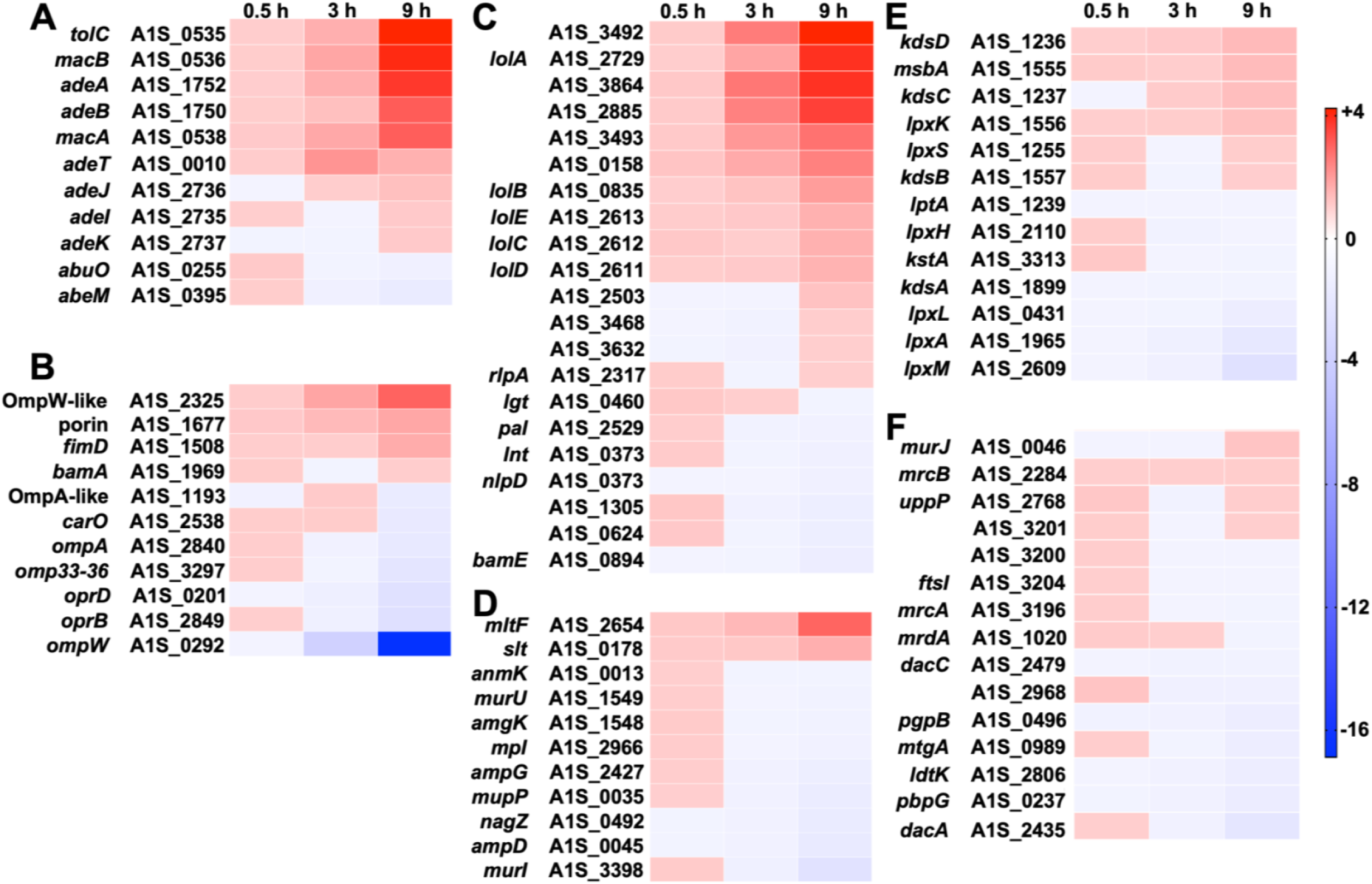
Differentially regulated genes in response to meropenem treatment in *A. baumannii*. Heat map showing the fold-change in genes expressed at 0.5, 3 and 9 h meropenem treatment relative to wild type ATCC 17978 (*P* <0.05). Pathways represented include genes associated with (A) efflux, (B) outer membrane porins, (C) outer membrane lipoproteins and their transporters, (D) peptidoglycan recycling, (E) LPS biosynthesis and (F) peptidoglycan biosynthesis.

Porins represent the major entryway for carbapenems such as meropenem to enter the periplasm^30^, where they inhibit transpeptidation to cross-link the stem peptides of adjacent PG strands. Decreased expression of many porin-associated genes was found in treated cultures relative to untreated (Fig 2B), suggesting the cell also limits meropenem entry by reducing porin gene expression in response to treatment. However, temporal expression of porin-associated genes was delayed relative to efflux, in general. Deletion of *carO* is associated with carbapenem resistance in *A. baumannii*^31^ and was found to be an influx channel for carbapenems^32^, while OprD has also been associated with clinical carbapenem resistance in *A. baumannii*^33^, suggesting reduced expression may strategically limit meropenem entry. Interestingly, the largest reduction in gene expression was associated with *ompW*, which encodes a predicted β-barrel protein (OmpW) that supports iron uptake^34^, but our understanding of its biological function or how it contributes to carbapenem resistance or tolerance is limited. Notably, in *Vibrio cholerae*, decreased iron uptake regulated by the VxrAB two-component system, promotes spheroplast recovery by reducing oxidative stress during β-lactam treatment^35, 36^.

As shown in separate *A. baumannii* transcriptional datasets in stress^37–39^, meropenem treatment also induces expression of genes encoding putative outer membrane lipoproteins and their transporters (LolA-D) (Fig 2C). Remodeling the outer membrane with lipoproteins presumably fortifies the envelope by providing structural rigidity, where inner leaflet outer membrane lipoproteins are covalently attached to the underlying PG network^1, 40^. Transcription of genes associated with PG remodeling were only slightly altered with the notable exception of two genes encoding putative lytic transglycosylases, including membrane-bound, MltF, and a soluble protein, Slt, which were both highly upregulated (Fig 2D). Lytic transglycosylases cleave *N*-acetylmuramic acid (Mur*N*Ac)-*N*-acetylglucosamine (Glc*N*Ac) bonds in PG to release soluble 1,6-anhydroMur*N*Ac-containing muropeptides. Muropeptides excised by lytic transglycosylases can be secreted into the environment or imported into the cytoplasm and catabolized via the PG recycling pathway^41^. 1,6-AnhydroMur*N*Ac-containing muropeptides that feed into PG recycling can act as a source of nutrients, but also can be re-incorporated into the PG network through *de novo* biosynthesis or in some bacteria can also act as signals to induce β-lactamase expression^42, 43^. Lastly, genes involved in lipooligosaccharide (LOS), and PG biosynthesis were slightly altered (Fig 2EF).

### Genes and pathways that contribute to *A. baumannii* fitness during meropenem treatment

While transcriptome sequencing analyses offer insight into the stress response, one limitation of RNA-sequencing is that differentially regulated genes oftentimes do not impact fitness due to redundancy or pleiotropic effects. Therefore, we also performed transposon-sequencing on *A. baumannii* strain ATCC 17978, as previously done^44, 45^, and compared recovered insertional mutants from meropenem treated and untreated cultures. The screen was answered by several novel factors, some of which are the subject of a separate study, but also revealed the importance for outer membrane integrity and PG maintenance. To validate our screen, we calculated survival in several mutants, including Δ*ompA*, Δ*lpxM*, Δ*pbpG* and Δ*ldtK (also known as elsL*^46^), in the presence and absence of meropenem (Fig 3A). All mutants showed a 2-to-3-fold log depletion in the mutants relative to wild type at 12 h and >5-fold log depletion at 24 h. These studies suggest that *A. baumannii* fitness during meropenem treatment is dependent on outer membrane and PG maintenance factors.

**Figure 3:**
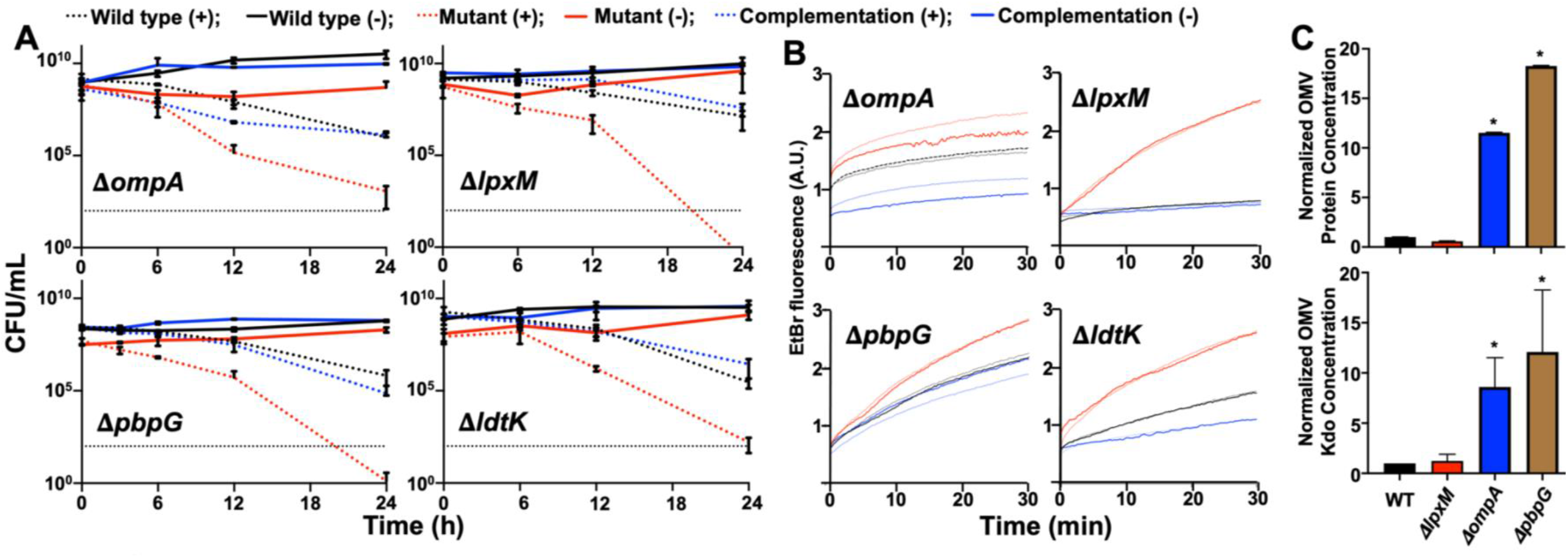
Genes encoding outer membrane integrity and peptidoglycan maintenance contribute to meropenem tolerance in *A. baumannii*. (A) Survival was calculated as CFU/mL over 24 h during meropenem treatment. Data were collected from two experiments in triplicate. Error bars represent the average of 3 technical replicates +/- standard deviation. (B) Permeability assays using ethidium bromide (EtBr) over 0.5 h. A.U.; arbitrary units. Lines shown depict the mean of three technical replicates. (C) Relative quantification of protein (top) and Kdo (bottom) concentrations of outer membrane vesicles (OMVs) in wild type (WT) and mutants. Each experiment was independently replicated three times, and one representative dataset was reported. Error bars indicate standard deviation. An asterisk indicates significant differences relative to the WT strain (*P* <0.05).

OmpA is a highly conserved monomeric β-barrel protein with a periplasmic domain that noncovalently attaches the outer membrane to the PG network^47^. It is a highly abundant protein in *A. baumannii*^48^ that couples with efflux pumps and can aid in export of antibacterial compounds from the periplasm^49, 50^. OmpA is known to stabilize the outer membrane; *ompA* deletion/disruption increases formation of outer membrane vesicles and permeability^51^. To test the hypothesis that *ompA* deletion perturbs the outer membrane to promote carbapenem entry in *A. baumannii*, we performed two assays, including permeability measurements (Fig 3B) and outer membrane vesicle quantification (Fig 3C). Consistent with other reports^51^, Δ*ompA* showed increased outer membrane vesicle formation and permeability to ethidium bromide, which is similar in size to meropenem. We also measured ethidium bromide influx in Δ*lpxM*, Δ*pbpG and* Δ*ldtK* (Fig 3B). Like Δ*ompA*, all isogenic mutations increased permeability relative to wild type and the respective complementation strain, which restored the permeability defect. Notably, meropenem treatment did not exacerbate permeability in wild type or any of the mutants (Fig 3B), suggesting it does not directly destabilize the outer membrane barrier function. Since we previously reported that Δ*ldtK* produces excess outer membrane vesicles^44^ and all of the mutants showed increased permeability, we also tested vesicle formation in Δ*lpxM* and Δ*pbpG* (Fig 3C). Unexpectedly, Δ*pbpG* produced excess outer membrane vesicles relative to wild type and all other mutants. In contrast, Δ*lpxM* did not.

Interestingly, Δ*lpxM* was the only strain that showed increased permeability but not hypervesiculation. LpxM catalyzes transfer of two lauroyl (C_12:0_) groups from an acyl carrier protein to the *R-*3*’*-and *R*-2-hydroxymyristate positions of lipid A during LOS biosynthesis^52^. Mutations that reduce LOS acylation are known to increase fluidity of the lipid bilayer and could also impact folding/function of outer membrane porins^53, 54^. Either/both mechanisms could increase entry of meropenem into the periplasmic space or disrupt efflux mechanisms that actively pump the compound out of the cell.

We also characterized the morphology of each mutant in growth (Fig S3). We found that relative to wild type, Δ*ompA* cells were chained and NADA incorporation was reduced (Fig S3A), suggesting that OmpA is required for proper function of PG enzymes (division proteins and LD-/DD-transpeptidases that incorporate NADA and/or increased carboxypeptidase activity). Δ*lpxM* and Δ*pbpG* showed increased NADA incorporation (Fig S3BC), which is consistent with increased outer membrane permeability. Δ*pbpG* cells were also clumped (FigS3C), suggesting the cells could not properly separate during division. As previously reported^44^, Δ*ldtK* showed rounded cells (Fig S3D). We also calculated the meropenem minimal inhibitory concentrations in each mutant, which did not significantly deviate from the parent strain (Fig S3E).

While the contribution of the *A. baumannii* outer membrane proteins OmpA and LpxM to gate carbapenem entry to promote antibiotic tolerance is straightforward, we were intrigued by genetic links to PG maintenance (i.e., *pbpG* and *ldtK*). While mutation of *pbpG* and *ldtK* impact outer membrane integrity (Fig 3), we also wanted to define their physiological role to determine specific pathways that contribute to meropenem tolerance.

### PBP7 is a DD-carboxypeptidase and endopeptidase that catalyzes formation of tetrapeptides

To define the activity of *A. baumannii* PBP7 (encoded by *pbpG*), we isolated PG from wild type and Δ*pbpG* in growth (Fig 4A; Table S1) and stasis (Fig 4B; Table S1). Muropeptides were generated by treatment with muramidase, separated by high-performance liquid chromatography and uncharacterized peaks were analyzed by tandem mass spectrometry, as done previously^37, 44, 55^. PG composition from Δ*pbpG* in growth showed accumulation of two muropeptide peaks that were not present in wild type (Fig 4A; Table S1). MS analysis showed these peaks were enriched with pentapeptides and were identified as disaccharide pentapeptide (Penta, neutral mass: 1012.19 amu; theoretical: 1012.45 amu) and bis-disaccharide tetrapentapeptide (TetraPenta, neutral mass: 1935.60 amu; theoretical: 1935.84 amu) (Fig 4C; Fig S4), suggesting the enriched muropeptide pools represent PBP7 substrates in growth. Δ*pbpG* PG in stasis had depleted D-amino acid-modified muropeptide pools, including TetraTri-D-Lys-and TetraTri-D-Arg-peptides, and reduced 3-3 crosslink formation (Fig 4BC; Table S1), consistent with PBP7 DD-carboxypeptidase and endopeptidase activity to form tetrapeptides, which are the most abundant peptides in the PG in *A. baumannii*^37, 44, 55^ and substrates of LD-transpeptidases. The periplasmic LD-transpeptidase, LdtJ, transfers D-amino acid to tetrapeptides and forms 3-3 crosslinks^44^. Therefore, it is likely that PBP7 provides at least some of the periplasmic substrates for LdtJ-dependent transpeptidase activity in stasis.

**Figure 4:**
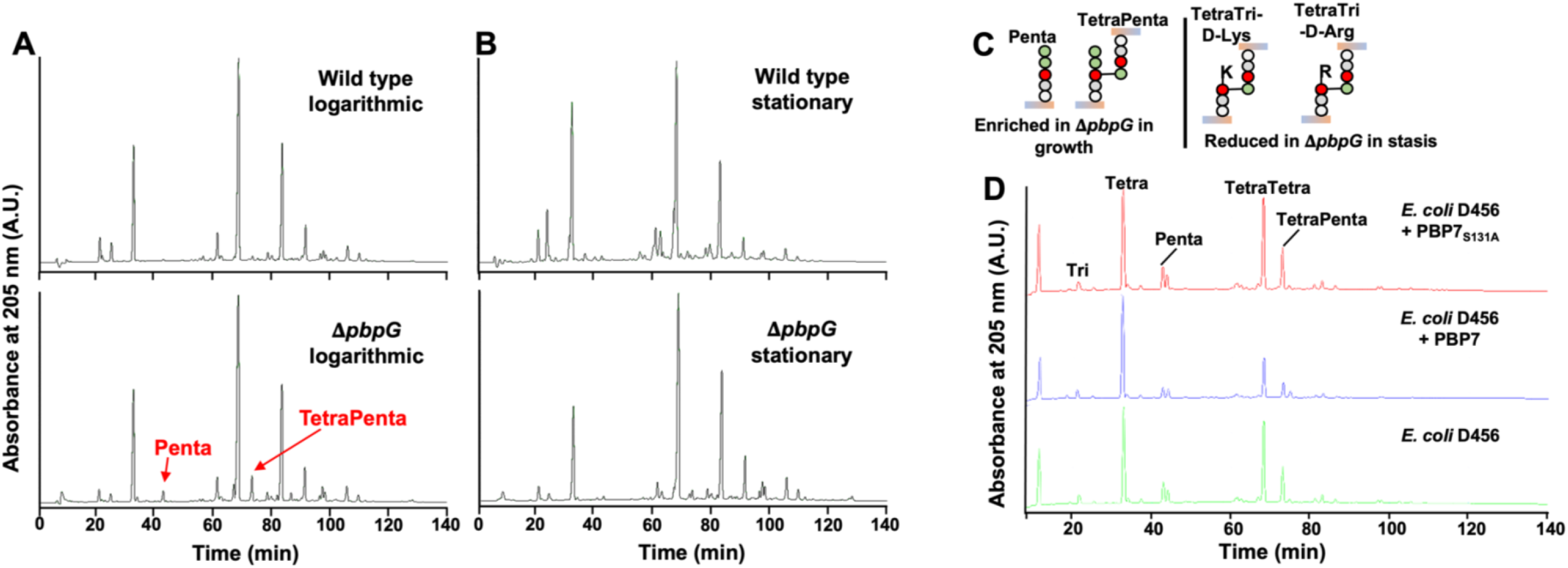
PBP7 is active against pentapeptides and DD-crosslinks. (A) PG isolated from wild type and **Δ***pbpG* in growth phase was analyzed by HPLC. The muropeptides Penta and TetraPenta were enriched in **Δ***pbpG.* (B) PG isolated from wild type and **Δ***pbpG* in stationary phase was analyzed by HPLC. TetraTri-D-Lys and TetraTri-D-Arg were depleted in **Δ***pbpG* relative to wild type. (C) Muropeptide structures are illustrated and were confirmed using MS/MS. (D) Recombinant PBP7 or the active-site mutant PBP7_S131A_ was incubated with PG isolated from *E. coli* D456 which contains Tetra, Penta, TetraTetra and TetraPenta as the main muropeptides. PBP7 was active against pentapeptides (DD-CPase) and cross-linked muropeptides (DD-EPase).

To test the enzymatic activity, we purified recombinant PBP7 (Fig S5A) and a predicted catalytically inactive version in which alanine replaces the active site serine (PBP7_S131A_). Purified proteins were incubated with PG from *E. coli* D456 (Fig 4D), a strain enriched with pentapeptides^56^ and analyzed as previously done^57^. PBP7 was active against penta-, tetratetra-and tetrapentapeptides, where each muropeptide was trimmed to the tetrapeptide-form relative to the no-enzyme control. As expected, PBP7_S131A_ did not show activity against any muropeptides. Together, these studies suggest that PBP7 not only hydrolyzes the bond between the terminal D-Ala residues, but also showed DD-endopeptidase activity and both activities enrich the periplasmic pool of monomeric tetrapeptides.

### LdtK is a cytoplasmic LD-carboxypeptidase active against tetrapeptides for PG recycling

During β-lactam treatment, autolysins (i.e., lytic transglycosylases) are activated^7, 58^, which increases the amount of PG turnover products with 1,6-anhydro-Mur*N*Ac residues. In *A. baumannii*, genes encoding the autolysins, MltF and Slt, were upregulated during meropenem treatment (Fig 2D), which likely increases periplasmic concentrations of TetraAnh, for cytoplasmic import. In *E. coli*, TetraAnh are substrates for the LD-carboxypeptidase LdcA, which trims tetrapeptides to tripeptides^59^ that are catabolized by the conserved enzymes NagZ^60, 61^ to generate 1,6-anhMur*N*Ac-tripeptide and the amidase AmpD^62, 63^ to form free tripeptides, which can be further broken down into individual amino acids and used as energy. Furthermore, Mpl^64^ can attach tripeptides to uridine diphosphate (UDP)-Mur*N*Ac to form UDP-Mur*N*Ac-tripeptide, an intermediate in the *de novo* PG biosynthesis pathway. However, no apparent LD-carboxypeptidase, orthologue to LdcA is encoded by *A. baumannii*.

LdtK was one potential LD-carboxypeptidase candidate for PG recycling because it encodes a putative LD-transpeptidase (YkuD) domain but does not encode a canonical secretion signal needed for export, suggesting it may be active in the cytoplasm. This observation coupled with a recent study showing that the *E. coli* YkuD homologue DpaA (also known as LdtF) is an amidase that hydrolyzes bonds formed by LD-transpeptidases^2, 65^, suggested that LdtK may indeed have LD-carboxypeptidase activity, which is essential for tripeptide formation in the recycling pathway.

First, we determined the subcellular localization of LdtK with a specific antibody that detects the native protein (Fig 5A). After fractionation of the subcellular compartments, we were only able to detect LdtK in the cytoplasmic fraction in growth and stasis, showing it is not exported to the periplasm.

**Figure 5:**
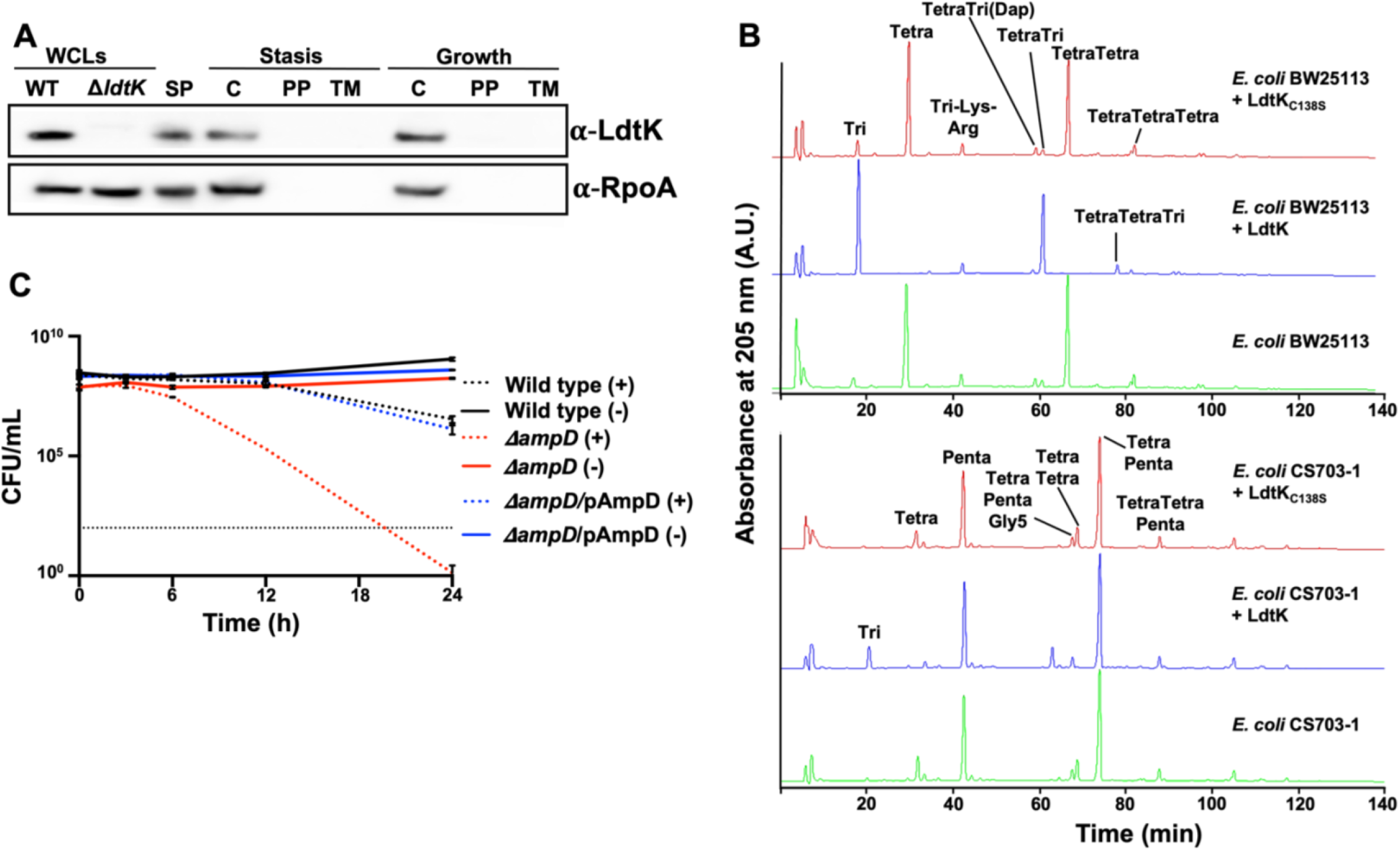
LdtK is active against tetrapeptides in Tetra and TetraTetra. (A) Western blot with **α**-LdtK and **α**-RpoA antisera. LdtK is 18.97 kDa, while RpoA is 37.27 kDa. WCL; whole-cell lysate, SP; spheroplast, C; cytoplasm, PP; periplasm, and TM; total membrane fractions. (B) Recombinant LdtK or the active-site mutant LdtK_C13S_ was incubated with PG isolated from *E. coli strain* BW25113 (top, tetrapeptide-rich) or strain CS703-1 (bottom, pentapeptide-rich). LdtK was active against tetrapeptides but not pentapeptides. (C) Meropenem tolerance assay in **Δ***ampD*, which encodes a well-conserved cytoplasmic enzyme required for PG recycling. Error bars represent the average of 3 technical replicates +/- standard deviation.

Since LdtK is cytoplasmic, we sought to determine if it was active on tetra- and/or pentapeptide substrates. We purified recombinant LdtK and the active-site mutant, LdtK_C138S_ (Fig S5B). Both enzymes were incubated with muropeptides obtained from tetrapeptide-rich PG from *E. coli* BW25113 or pentapeptide-rich PG from *E. coli* CS703-1^57^ (Fig 5B). LdtK showed activity against tetrapeptides but not pentapeptides. The muropeptide profile showed the formation of disaccharide tripeptide and bis-disaccharide tetratripeptide, showing that LdtK cleaves the bond between the L-centre of *m*DAP and the terminal D-Ala in tetrapeptides, characteristic of LD-carboxypeptidase activity. Our data indicate that PBP7 trims pentapeptides and cleaves crosslinked peptides into tetrapeptides in the periplasmic PG network. Once 1,6-anhydro-Mur*N*Ac-containing muropeptides are released from the PG network by MltF, Slt or other lytic transglycosylases, they are transported into the cytoplasm and into tripeptides by LdtK in the PG recycling pathway.

To further confirm if the cytoplasmic PG pathway contributes to meropenem tolerance in *A. baumannii*, we made an isogenic *ampD* mutant, which encodes *N*-acetylmuramyl-L-alanine amidase that releases the tripeptide from anhMur*N*Ac^63^. Like Δ*pbpG* and Δ*ldtK*, Δ*ampD* was also rapidly killed when treated with meropenem relative to wild type and the respective complementation strain (Fig 5C). Together, these studies strongly suggest that the PG recycling pathway contributes to meropenem tolerance in *A. baumannii*. Furthermore, the formation of cytoplasmic tripeptides or tetrapeptides appears to contribute to meropenem tolerance. Combinatorial therapies that inhibit enzymes in both PG biosynthesis and recycling could provide an alternative treatment strategy.

## Discussion

Many susceptible Gram-negative pathogens tolerate treatment with bactericidal antibiotics such as carbapenem β-lactams, but the molecular factors that underlie cell survival are not understood. Significant cell envelope damage is observed during treatment, characterized morphologically by cell wall-depleted spheroplasts^17–21^. Here, we show meropenem treatment induces spheroplast formation in *A. baumannii*, and that cell growth resumes upon removal of the antibiotic. Transcriptome sequencing analysis suggests *A. baumannii* responds to treatment by fortifying the structural integrity of the cell envelope through increased outer membrane lipoprotein and transporter gene expression and by inducing autolysins, which likely physically reinforce the envelope by providing stiffness and remodel the PG network, respectively. Treated cells also appear to limit intracellular meropenem concentrations through induced expression of efflux-associated genes and downregulation of porin genes, which both reduce periplasmic concentrations by actively pumping the antibiotic out of the cell and by limiting entry, respectively. A separate genetic (transposon) screen to identify fitness determinants, showed factors required for high level meropenem tolerance include genes that contribute to outer membrane permeability (*lpxM*, *ompA*, *pbpG* and *ldtK*) and cell envelope stability (*ompA*, *pbpG* and *ldtK*). Furthermore, genes in the cytoplasmic PG recycling pathway, *ldtK* and *ampD*, also answered the screen. Together the transcriptomics and genetic screen suggested factors that maintain cell envelope homeostasis through integrity maintenance of the outer membrane and PG network contribute to meropenem tolerance in *A. baumannii*.

While we showed several tolerance factors are transcriptionally regulated, we do not know what transcription factors are involved. A previous study found that PhoPQ-dependent outer membrane modifications promoted survival in cell wall-deficient spheroplasts^20^, presumably by fortifying the outer membrane to counter large loads of turgor pressure typically absorbed by the cell wall. Specifically, PhoPQ was activated in response to meropenem treatment. *A. baumannii* does not encode PhoPQ, but analogous mechanisms likely to contribute to cell envelope homeostasis to counter the turgor when the cell wall is compromised. One mechanism might include fortification of the cell envelope with lipoproteins, which occurs in stress in *A. baumannii*^37–39, 66^; however, the underlying protective mechanism is not understood. Landmark studies in cell envelope mechanics have shown that outer membrane lipoprotein attachment to the PG impact cell envelope mechanics by increasing the load-bearing capacity of the cell envelope^1, 40^. Outer membrane lipoproteins, specifically those that interact with the underlying PG network, increase outer membrane stiffness, which likely counterbalances the internal turgor. When the cell wall is perturbed during meropenem treatment, small fluctuations in turgor may be sufficient to induce lysis when lipoprotein-mediated attachment is absent. More studies are needed to tease apart the contribution of specific lipoproteins and how they contribute to cell envelope mechanics in stress. Furthermore, noncovalent attachments between the outer membrane and PG network via OmpA and hyperacylation of lipid A via LpxM may also increase the mechanical load-bearing capacity of the outer membrane to maintain envelope homeostasis when the cell wall is defective. It is also possible that disruption of OmpA or LpxM has pleiotropic effects that reduce the barrier function to gate meropenem entry.

Unexpectedly, our data suggest that PG maintenance enzymes contribute to *A. baumannii* survival during meropenem treatment. Tetrapeptides represent the most abundant PG stem peptides in *A. baumannii*. They are formed, in part, by the DD-carboxypeptidase and endopeptidase activity of PBP7 from pentapeptides and DD-crosslinked muropeptides, respectively (Fig 4A). Tetrapeptides are substrates for LD-transpeptidase that form a small amount of 3-3 crosslinks in *A. baumannii*, but are needed to effectively repair PG defects in stressed *E. coli* cells^2^. Furthermore, tetrapeptides are also necessary for LD-transpeptidase-dependent covalent attachment of Braun’s lipoprotein (Lpp) to *meso*-DAP residues in PG^67^, which also fortifies the envelope^1^. Our data indicate that without PBP7, LdtJ-dependent 3-3 crosslink formation is reduced (Fig 4A), however less than 3% of all muropeptides contain 3-3 crosslinks and their contribution to PG integrity maintenance remain unclear.

The lytic transglycosylases MltF and Slt were induced in meropenem treatment (Fig 2), consistent with activation of autolysins in response to penicillin-binding protein inhibition during β-lactam treatment^7, 17^. Lytic transglycosylase proteins cleave the glycosidic linkage between disaccharide subunits within the PG strands and perform an intramolecular transglycosylation in Mur*N*Ac to release soluble 1,6-anhydroMur*N*Ac-containing muropeptides, which can be imported into the cytoplasm. The main turnover product, TetraAnh is transported into the cytoplasm by AmpG where they provide substrates for LdtK-dependent LD-carboxypeptidase activity to form TriAnh. Like other members of the YkuD family, LdtK retains preference for tetrapeptide substrates (Fig 5B) but represents the first known YkuD-containing enzyme that lacks a signal sequence and is active in the cytoplasm. LdtK is the second member of the YkuD family, after DpaA/LdtF that has a major role in cleaving amide bonds rather than generating them. Notably, the requirement for LdtK in meropenem tolerance is like how cells depend on outer membrane integrity maintenance (OmpA, LpxM, PBP7) during spheroplast formation. Lastly, LdtK (and LdtJ) were shown to be essential for *A. baumannii* survival without LOS^44^, suggesting that PG recycling and modification of tetrapeptides are a general response to counter cell envelope stress in *A. baumannii*. It is also possible that cytosolic accumulation of tetrapeptides creates another problem in cells that are already sick and cannot be tolerated.

Hydrolysis of TriAnh by the dedicated enzymes NagZ and AmpD, which lead to the formation of anhMur*N*Ac-tripeptide, 1,6-anhMur*N*Ac and tripeptides, respectively, could be degraded into individual amino acids for utilization as nutrient or energy sources^42, 68, 69^ to promote survival during tolerance. It is reasonable to expect the cell requires some nutrients during tolerance, and this pathway could provide energy to support basal metabolic processes. Alternatively, Mpl could ligate tripeptides to UDP-Mur*N*Ac in the recycling pathway^64, 70^. UPD-Mur*N*Ac-tripeptide is an intermediate in the *de novo* PG synthesis pathway^71–73^; however, it is not obvious how *de novo* PG synthesis via recycling would benefit the bacterium during treatment because periplasmic PBPs and LD-transpeptidases, which are required for crosslinking, are inhibited by meropenem. Another possibility is that accumulation of cytoplasmic 1,6-anhydroMur*N*Ac-containing muropeptides provide signals to induce β-lactamase expression, which could localize in the periplasm to reduce meropenem concentrations to survivable levels. Two mechanisms have been characterized in Gram-negative bacteria, including the AmpG-AmpR pathway and the BlrAB two-component system, which both induce β-lactamase expression in response to muropeptide concentrations^43^. Many genes in *A. baumannii* have not yet been characterized. Signaling pathways and potentially carbapenemases could be induced in response to 1,6-anhydroMur*N*Ac-containing muropeptide accumulation to promote meropenem degradation. A more detailed analysis is needed to characterize the PG recycling tolerance mechanism, which will inform more effective treatment strategies to combat *A. baumannii* infections. Furthermore, our studies also show that disruption of PG maintenance enzymes (i.e., PBP7, LdtK) compromised outer membrane integrity. It is also possible that outer membrane perturbations in these mutants induce unchecked antibiotic entry to impact fitness during meropenem treatment. Notably, regulatory links between the outer membrane and PG maintenance are not well-understood in *A. baumannii*.

## Materials and Methods

### Bacterial strains and growth

All strains and plasmids used in this study are listed in Table S2 in the supplementary material. Primers are listed in Table S3. All *A. baumannii* strains were grown aerobically from freezer stocks on Luria-Bertani (LB) agar at 37° C. Antibiotics were used at the following concentrations unless noted otherwise: 25 mg/L kanamycin, 10 mg/L meropenem, 10 mg/L tetracycline.

### Construction of genetic mutants

*A. baumannii pbpG, ampD, ompA* mutants were constructed as described previously^44, 52^, using the recombination-mediated genetic engineering (recombineering) method^52^. Briefly, a kanamycin resistance cassette flanked by FLP recombination target (FRT) sites was PCR amplified from the pKD4 plasmid using primers containing 125-bp flanking regions of homology to the gene of interest. The resulting linear PCR product was then transformed via electroporation into *A. baumannii* strains ATCC 17978 expressing pREC*_Ab_* (pAT03). Transformants were recovered in Luria broth and plated on LB agar supplemented with 7.5 mg/L kanamycin. All genetic mutants were confirmed by PCR.

Following isolation of genetic mutants, the pMMB67EH::REC*_Ab_* Tet^r^ plasmid was removed as described previously^52^. Isolated mutants were grown on LB agar supplemented with 2 mM nickel (II) chloride (NiCl_2_) and replica plated on LB agar supplemented with kanamycin or tetracycline. Loss of pMMB67EH::REC*_Ab_* Tet^r^ plasmid in mutants susceptible to tetracycline and resistant to kanamycin were confirmed using PCR. To excise chromosomal insertion of the kanamycin resistance cassette, cured mutants were transformed with pMMB67EH carrying the FLP recombinase (pAT08) and plated on LB agar supplemented with tetracycline and 2mM Isopropyl β-d-1-thiogalactopyranoside (IPTG) to induce expression of FLP recombinase. Successful excision of the kanamycin resistance cassette was confirmed using PCR.

pPBP7 was constructed by amplifying the *pbpG* (A1S_0237) coding sequence (encoding PBP7) with 200-bp upstream and downstream flanking regions was amplified from *A. baumannii* ATCC 17978 chromosomal DNA (cDNA) and cloned into XhoI and KpnI restriction sites in the pABBRkn^R^ plasmid. The resulting pPBP7 plasmid was transformed into *A. baumannii* ATCC 17978 Δ*pbpG* background for complementation using the native promoter.

AmpD and OmpA complementation vectors were constructed similarly with slight alterations. The *ampD* (A1S_0045) and *ompA* (A1S_2840) coding sequences were amplified from *A. baumannii* ATCC 17978 cDNA and cloned into BamHI and SalI restriction sites in the pMMB67EHkn^R^ plasmid. The resulting pAmpD and pOmpA plasmids were transformed into the respective mutant and induced with 2 mM IPTG for complementation.

### Fluorescent NADA staining

Overnight cultures were grown with shaking at 37° C in 5 mL of BHI (BD Difco Bacto Brain Heart Infusion) broth. The following day, cultures were back diluted at 1:10 in fresh BHI media (total volume 5 mL) containing without or with meropenem. 2 μL of 10 mM NBD-(linezolid-7-nitrobenz-2-oxa-1,3-diazol-4-yl)-amino-D-alanine (NADA) (Thermo Fisher) was added to each tube and incubated at 37° C. At noted time points 6, 12, and 24 h, cultures (5 mL) were washed twice in BHI broth and fixed with phosphate-buffered saline containing a (1:10) solution of 16% paraformaldehyde. For spheroplast recovery, 12 h treated cultures were washed 3 times in BHI to remove the excess meropenem and resuspended in fresh BHI. 10 mM NADA was added and incubated for 12 h at 37° C before fixing the cells for microscopy.

### Microscopy

Paraformaldehyde-fixed cells were immobilized on 1.5% agarose pads and imaged using an inverted Nikon Eclipse Ti-2 widefield epifluorescence microscope equipped with a Photometrics Prime 95B camera and a Plan Apo 100x 1.45-numerical-aperture lens objective. Phase-contrast and fluorescence images were collected with NIS Elements software. Green fluorescence images were taken using a Sola LED light engine and filter cube with 632/60 or 535/50 emission filters.

### Image analysis

Microscopy images were processed and pseudo colored with ImageJ Fiji^74^. A cyan lookup table was applied to NADA images. Cells shape (length, area, width and fluorescence intensities) were quantified in MicrobeJ^75^ and data were plotted in Prism 9 (GraphPad 9.2.0). Each experiment and independently replicated three times, one representative data was reported in the quantification and one representative image was included in the figure.

### RNA-sequencing

Transcriptome sequencing analysis was performed as described previously with modification^37^. Briefly, the Direct-Zol RNA MiniPrep kit (Zymo Research) was used to extract total RNA from *A. baumannii* ATCC 17978 cultures either treated with meropenem or an equivalent volume of water as blank at 0.5, 3, and 9 h at 37° C in triplicate. Turbo DNA-free DNA removal kit (Invitrogen) was used to remove genomic DNA contamination. DNase-depleted RNA was sent to the Microbial Genome Sequencing Center (MiGS) for Illumina NextSeq 550 sequencing. CLC genomic workbench software (Qiagen) was used to align the resulting sequencing data to the *A. baumannii* ATCC 17978 genome annotations and determine the RPKM expression values and the weighted proportions fold change of expression values between meropenem treated and treated samples. Baggerley’s test on proportions was used to generate a false discovery rate adjusted *P*-value. The weighted proportions fold change of expression values between samples was used to generate pathway-specific heatmaps in Prism 9. The sequencing data has been deposited in the National Center for biotechnology’s Gene Expression Omnibus.

### Transposon insertion sequencing

Transposon sequencing was performed as described previously^37, 44, 45, 76^. Briefly, pJNW684 was conjugated into wild-type *A. baumannii* strain ATCC 17978 to generate a library of ∼400,000 mutants. The transposon mutant library was pooled and screened for survival with and without meropenem treatment at 6 h at 37° C. Genomic DNA (gDNA) from meropenem treated and untreated cultures was isolated, sheared and transposon junctions were amplified and sequenced. Frequency of transposon insertions was compared between meropenem treated and untreated conditions to determine fitness determinants that contribute to carbapenem tolerance in *A. baumannii*.

### Time-dependent killing assays

Meropenem killing experiments were performed as previously described with slight alteration^77^. Wild type, mutant and complementation strains were grown overnight in Luria broth at 37° C. The following day, overnight cultures were back diluted 1:10 in fresh, pre-warmed BHI broth containing meropenem or an equivalent volume of water. Diluted BHI cultures were then incubated at 37° C. At 0, 3, 6, 12, and 24 h, each sample was diluted 4-fold in blank BHI, and the optical density (OD_600_) was measured. At each time point, cells were serially diluted 10-fold in fresh BHI broth and either 5 μL of each serial dilution was spot-plated, or 100 μL of each dilution was plated on LB agar. Spot-plates were imaged and CFUs were calculated after 24 h at 37° C. Each experiment was independently replicated three times, and one representative dataset was reported.

### Construction of PBP7 and LdtK active-site mutants

Site-directed mutagenesis was performed, as previously described with *ldtK (*A1S_2806)^44^. Briefly, the *pbpG* coding sequence was amplified from *A. baumannii* ATCC 17978 cDNA, cloned into the BamHI restriction site in pUC19 and transformed into *E. coli* C2987 chemically competent cells (New England Biolabs, Inc). pUC19::PBP7 was used as a template for Pfu-mediated deletion mutagenesis. DpnI-digested PCR reactions were transformed into *E. coli* C2987 chemically competent cells and plated on LB agar supplemented with 75 mg/L carbenicillin. All mutants were confirmed by PCR and Sanger sequencing.

### Construction of PBP7 and LdtK overexpression strains

*pbpG* and *ldtK* coding sequences were amplified from *A. baumannii* ATCC 17978 cDNA and *pbpG*_S131A_ and *ldtK*_C138S_ were amplified from pUC19::*pbpG*_S131A_ and pUC19::*ldtK*_C138S_ plasmid DNA using primers containing his_8X_-tag sequence. Amplicons were cloned into NdeI and BamHI restriction sites in pT7-7Kn and transformed into *E. coli* C2987 chemically competent cells, resulting pT7-7Kn::*pbpG*, pT7-7Kn::*pbpG*_S131A_, pT7-7Kn::*ldtK* and pT7-7Kn::*ldtK*_C138S_. Constructs were confirmed using Sanger sequencing and transformed into chemically competent *E. coli* C2527 (BL-21) (New England Biolabs, Inc) for purification, expression and western blotting.

### Purification of recombinant PBP7 and LdtK

BL21 cells containing carrying pT7-7Kn::*pbpG*, pT7-7Kn::*pbpG*_S131A_, pT7-7Kn::*ldtK* and pT7-7Kn::*ldtK*_C138S_ were grown in 500 mL Luria broth and 1 mM IPTG at 37° C for 7 h. Cells were collected and washed in cold 1x Phosphate-buffered saline (PBS), pelleted and the supernatant was removed. The dry pellet was frozen at −80° C overnight. The pellet was thawed on ice and resuspended in 20 mL lysis buffer (20 mM Tris, 300 mM NaCl, 10 mM imidazole; pH 8). Samples were sonicated for 20 s on and off for 10 min at 60% amplitude (Qsonica Q125 Sonicator). Cells were centrifuged at 20,000 x g for 0.5 h at 4°C. Supernatant was incubated with lysis buffer washed HisPur Ni-NTA Resin (Thermo Scientific) on a rotator for 2 h at 4°C. Sample was added to a 10 mL protein purification column containing a porous polyethylene disk (Thermo Scientific) and allowed to gravity drip. The column was washed 3x with 20 mL lysis buffer and increasing concentrations of additional imidazole at each wash (0 mM, 15 mM, and 30 mM). 500 μl of elution buffer (20 mM Tris, 300 mM NaCl, 250 mM imidazole; pH 8) was incubated with the column for 5 min then gravity eluted 9 times. The elution fractions containing protein, as determined by a protein gel, were injected into a 10-mW dialysis cassette (Thermo Scientific) and dialyzed over-night in dialysis buffer (10 mM Tris, 50 mM KCl, 0.1 mM EDTA, 5% glycerol; pH 8) at 4°C. Purified protein was collected and verified using western blot with an anti-his antibody.

### Isolation of outer membrane vesicles

Outer membrane vesicles were isolated as described previously^44^. Briefly, overnight cultures were back-diluted to OD_600_ 0.01 in 100 mL Luria broth and grown to stationary phase at 37° C. Cultures were then pelleted at 5000 x g for 15 min at room temperature and the supernatant was filtered through a 0.45 mm bottle-top filter. Filtered supernatant was ultracentrifuged (Sorvall WX 80+ ultracentrifuge with AH-629 swinging bucket rotor) at 151,243 x g for 1h at 4° C. Following final ultracentrifugation, outer membrane vesicle pellet was resuspended in 500 mL cold membrane vesicle buffer (50 mM Tris, 5 mM NaCl, 1 mM MgSO_4_; pH 7.5). Outer membrane vesicles were repeated three times in duplicate, one representative data set was reported.

### Quantification of total outer membrane vesicle proteins

Bradford assay was used to determine outer membrane vesicle protein concentration, as previously described^44^. To generate a standard curve, bovine serum albumin (BSA) was diluted 0 to 20 mg/mL in Pierce Coomassie Plus assay reagent (ThermoFisher) to a final volume of 1 mL. Outer membrane vesicles were diluted 2, 5, 10, 15, 20 μL in reagent to a final volume of 1 mL. A microplate spectrophotometer (Fisherbrand AccuSkan) was used to measure the absorbance (OD_595_) of standard and samples in a 96-well plate (BrandTech). Protein concentrations were determined by comparing the optical densities of samples to the standard curve plotted in Microsoft Excel and final quantifications were graphed in GraphPad Prism 9. Experiments were reproduced three times from each outer membrane vesicle isolation, and one representative data set was reported.

### Quantification of outer membrane vesicle 3-deoxy-**D**-*manno*-oct-2-ulosonic acid (Kdo) concentrations

Kdo assays were carried out as described previously^44, 78^. For the standard curve, Kdo standard (Sigma) was diluted 0 to 128 μg/mL in 50 μL of DI water. 50 μL of 0.5 M sulphuric acid (H_2_SO_4_) was added to 50 μL of isolated outer membrane vesicles and freshly prepared 50 μL dilutions of the Kdo standard. Outer membrane versicles in 0.5 M H_2_SO_4_ were boiled for 8 min to release the Kdo sugars. Samples were allowed to cool for 10 min at room temperature. 50 μL of 0.1 M periodic acid was added to outer membrane vesicles and Kdo standards and incubated at room temperature for 10 min. Following incubation, 200 μL of 0.2 M sodium arsenite in 0.5 M hydrochloric acid (HCl) was added to outer membrane vesicles and Kdo standards followed by 800 μL of 0.6% freshly prepared thiobarbituric acid (TBA). All samples were boiled for 10 min and allowed to cool at room temperate for 30-40 min. Prior to optical density measurements, purified Kdo was extracted using *n*-butanol equilibrated with 0.5M HCl. Optical density was measured at OD_552_ and OD_509_ (Fisherbrand AccuSkan microplate spectrophotometer) in disposable polystyrene cuvettes (Fisherbrand). A linear Kdo standard curve was generated by subtracting OD_552_ measurements from OD_509_ measurements in Microsoft Excel and final quantifications were graphed in GraphPad Prism 9. Experiments were reproduced three times from each outer membrane vesicle isolation, and one representative data set was reported.

### Ethidium bromide permeability assay

Permeability assays were done as previously described^79^, with slight modifications. Briefly, overnight cultures were grown in 5 mL BHI medium, normalized and back diluted (1:10) in BHI with and without meropenem. Cultures were withdrawn at 0, 6 and 12 h and washed 3 times with PBS and normalized based on OD_600_. 180 mL of the cultures was added to 96 well black plate and 6μM EtBr was added immediately before fluorescence measurements. The relative fluorescence unit was analyzed using synergy multi-mode plate reader (530 nm excitation filter, 590 nm emission filter and 570 nm dichroic mirror). The temperature was adjusted to 25° C and read at 15 s intervals for 0.5 h. Assays were repeated three times in triplicate, one representative data set was reported. The mean RFU for each sample was calculated and plotted by Prism 9. Experiments were reproduced three times, and one representative data set was reported.

### PG isolation

Biological replicates were grown to mid-logarithmic or stationary phase in 400 mL of Luria broth. Cells were centrifuged (Avanti JXN-26 Beckman Coulter Centrifuge, Beckman Coulter JA-10 rotor) at 7,000 x g for 0.5 h at 4°C, resuspended in chilled 6 mL PBS and lysed via drop-wise addition to boiling 8% sodium dodecyl sulfate (SDS). PG was further purified as previously described^80^. Briefly, muropeptides were cleaved from PG by Cellosyl muramidase (Hoechst, Frankfurt am Main, Germany), reduced with sodium borohydride and separated on a 250-by 4.6-mm 3-μm Prontosil 120-3-C_18_ AQ reversed-phase column (Bischoff, Leonberg, Germany). The eluted muropeptides were detected by absorbance at 205 nm. Eluted peaks were designated based on known published chromatograms^37, 44, 55^; new peaks analyzed by MS/MS, as previously done^44^.

### Activity Assays

PBP7 activity assays were carried out in a final volume of 50 µl containing 20 mM Hepes pH 5.0, 6.0 or 7.5, 50 mM NaCl and 2 µM PBP7 or PBP7_S131A_. PG from *E. coli* D456 was added, and the reaction mixture was incubated at 37° C for 16 h. The reaction was stopped by boiling the samples for 10 min. The reaction was reduced with sodium borohydride and acidified to pH 4.0-4.5. *E. coli* D456 PG at pH 5.0 buffer conditions served as control. Muropeptides were analyses as previously described^80^.

LdtK activity assays were carried out in a final volume of 50 µl containing 20 mM NaP pH 5.0 and 10 µM LdtK or LdtK_C138S_. PG from *E. coli* BW25113 (WT, tetra-muropeptide-rich) or CS703-1 (multiple mutations in penicillin-binding proteins, penta-muropeptide-rich) was added, and the reaction mixture was incubated at 37° C for 4 h. The reaction was stopped by boiling the samples for 10 min. Muropeptides were reduced with sodium borohydride. Muropeptides were analyses as previously described^80^.

### Protein Localization

Cells were grown to mid-logarithmic or stationary phase and normalized to a density of OD_600_ 0.75 in 20 mL. Cultures were washed twice with chilled 1x PBS + 0.1% gelatin (PBSG) and resuspended in chilled 2 mL PBSG containing 2mg/mL Polymyxin B sulfate (MilliporeSigma) then agitated for 0.5 h at 4° C. Spheroplasts pelleted at 20,000 x g for 0.5 h at 4° C. The remaining supernatant was centrifuged at 20,000 x g for 0.5 h at 4° C. Supernatant was collected and saved as the periplasmic fraction. Previously pelleted spheroplasts were resuspended in 1 mL 10mM 4-(2-hydroxyethyl)-1-piperazineethanesulfonic acid (HEPES) buffer solution (Gibco) and 100 μl were collected as whole spheroplasts. Remaining spheroplasts were sonicated 15 seconds on and 15 seconds off 10 times at 60% amplitude (Qsonica Q125 sonicator). Lysed spheroplasts were pelleted at 16,000 x g for 0.5 h at 4°C. Supernatant was centrifuged another 0.5 h for 16,000 x g at 4°C. Insoluble pellet was saved as total membrane fraction. Soluble supernatant was saved as the cytoplasmic fraction. Experiments were reproduced three times, and one representative data set was reported.

### Immunoblots

All western blot analysis was performed using 4-12% Bis-Tris 10-well protein gels (Invitrogen) and NuPage MES SDS running buffer (Novex). Gels were transferred with NuPage transfer buffer (Novex) to 0.45 μm polyvinylidene difluoride (PVDF) (Amersham Hybond) membranes. All blots were blocked in 5% milk and 1x tris-buffered saline (TBS) for 2 h. Primary rabbit antisera, anti-LdtK and anti-RpoA were used at 1:750 dilution. Anti-rabbit horseradish peroxidase (HRP) secondary antibody was used at 1:10,000 (Thermo Fisher Scientific). Primary mouse antisera, 5x-His mouse anti-tag was used at 1:500 (Invitrogen). Anti-mouse HRP secondary antibody was used at 1:10,000 (Invitrogen). SuperSignal West Pico Plus (Thermo Fisher Scientific) was applied to detect relative protein concentrations.

For localization assays: whole cell lysate, whole spheroplasts and membrane fractions were mixed with 1x loading dye containing 4% 2-Mercaptoethanol (Fisher Chemical) and boiled for 10 min. 10 μl of each sample was used. 132 μl of the periplasmic or cytoplasmic fractions were added to 66 μl of 3x loading buffer containing 4% 2-Mercaptoethanol (Fisher Chemical) and boiled for 10 min. 60 μl of each sample was used. Each sample was loaded into 4-12% Bis-Tris 10-well protein gels (Invitrogen) for immunoblotting.

For protein purification: 1 μg of purified protein was combined with 3x loading buffer containing 4% 2-Mercaptoethanol and boiled for 10 min. The sample was loaded into 4-12% Bis-Tris 10-well protein gel (Invitrogen) for immunoblotting.

### Polyclonal antibody generation

Peptide fragments of LdtK and RpoA was used to generate two rabbit polyclonal antibodies against by Life Technologies Corporation (Grand Island, NY). Collected serum was tested for LdtK and RpoA reactivity in an enzyme-linked immunosorbent assay (ELISA) with peptide fragments and via western blot against whole-cell lysates.

### Minimal inhibitory concentrations (MICs)

MICs were determined using the Broth Microdilution (BMD) method, as previously outlined ^81^. Overnight cultures were back diluted to OD_600_ 0.01 and 100 μL of cells was added to each well of a 96-well round-bottom polypropylene plate (Grenier Bio-One). Meropenem diluted in water was serially diluted and 150 μL of each meropenem serial dilution was also added to each well. Plates were incubated overnight at 37°C and growth was measured by reading OD_600_ after 24 h of incubation. The lowest concentration of meropenem at which no bacterial growth was observed was determined to be the MIC. Assays were repeated three times in triplicate, one representative data set was reported.

### Statistical Analysis

Tests for significance in cell morphology, fluorescence intensity, outer membrane vesicle production were conducted using the Student *t*-test (two-tailed distribution with two-sample, equal variance calculations). Statistically significant differences between relevant strains possessed *P* <0.05.

