## Supplemental material for "Peptidoglycan recycling contributes to outer membrane integrity and carbapenem tolerance in *Acinetobacter baumannii*"

**Supplemental material for 'Peptidoglycan recycling promotes outer membrane integrity and carbapenem tolerance in *Acinetobacter baumannii*'**

Nowrosh Islam, Misha I. Kazi, Katie N. Kang, Jacob Biboy, Joe Gray, Feroz Ahmed, Richard Schargel, Cara C. Boutte, Tobias Dörr, Waldemar Vollmer, Joseph M. Boll

**Supplementary figures**

**Figure S1**

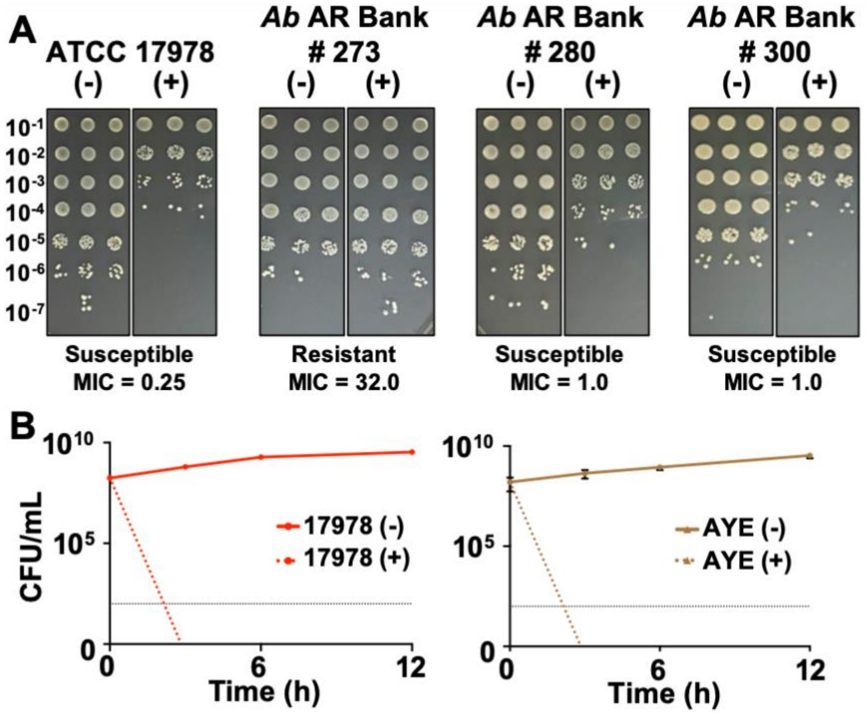

**Figure S1: Tolerance in clinical *A. baumannii* isolates.** (A) Dilution spot assays of *A. baumannii* strain ATCC 17978 and three recent clinical isolates, including a resistant (*A. baumannii* AR Bank # 273) and two meropenem susceptible (*A. baumannii* AR Bank # 280 and # 300) strains for comparison. The calculated meropenem minimal inhibitory concentration (MIC) is indicated below each image. (B) Survival (CFU/mL) of *A. baumannii* strains ATCC 17978 and AYE in logarithmic phase cultures was calculated over 12 h during meropenem treatment. Each experiment was independently replicated two times, and one representative data set was reported. Dotted black line indicated level of detection. Error bars represent the average of 3 technical replicates +/- standard deviation

**Figure S2**

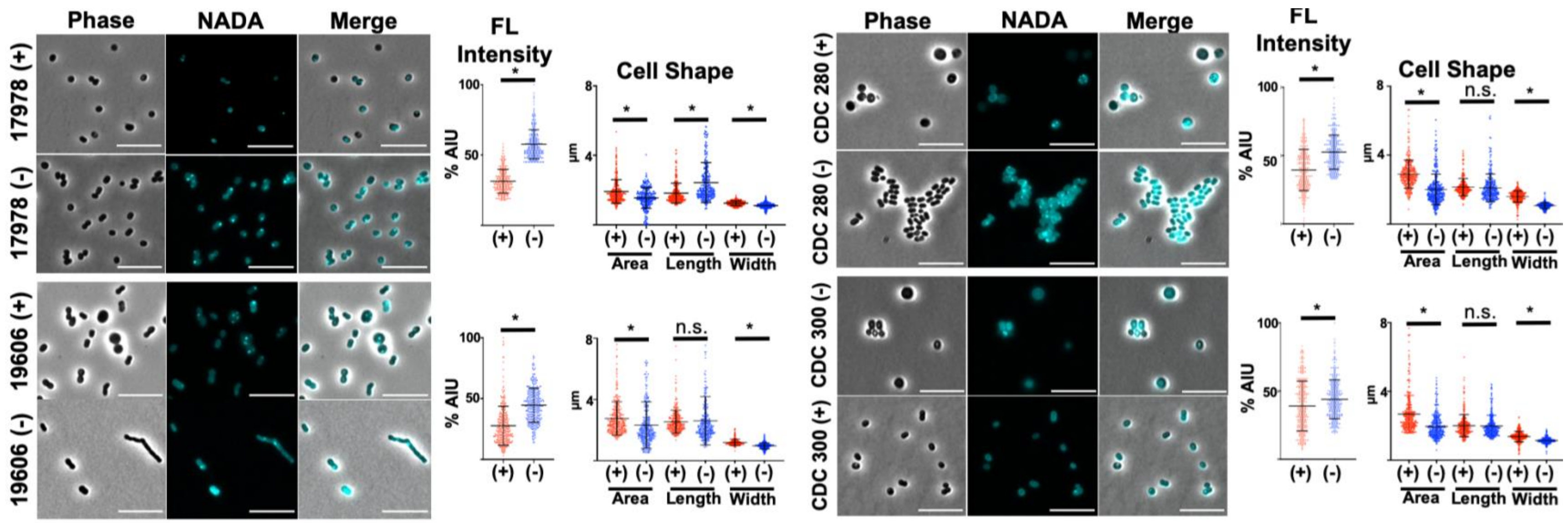

**Figure S2: Spheroplast formation in *A. baumannii* isolates after 12 h meropenem treatment.** Phase and fluorescence microscopy of treated (+) or untreated (-) *A. baumannii* strains after 12 h. Scale bar is 10 µm. To the right of each image are fluorescence (FL) intensity quantifications reported in percent arbitrary intensity units (AIU) in treated (red, +) vs. untreated (blue, -) and cell shape quantifications including, area (A), length (L) and width (W) (*n*= 300). Significance was determined using an unpaired t-test ($P$ <0.05) in treated vs. untreated. Error bars indicate standard deviation relative to the mean. An asterisk indicates significant differences ($P$ <0.05); n.s., not significant.

**Figure S3**

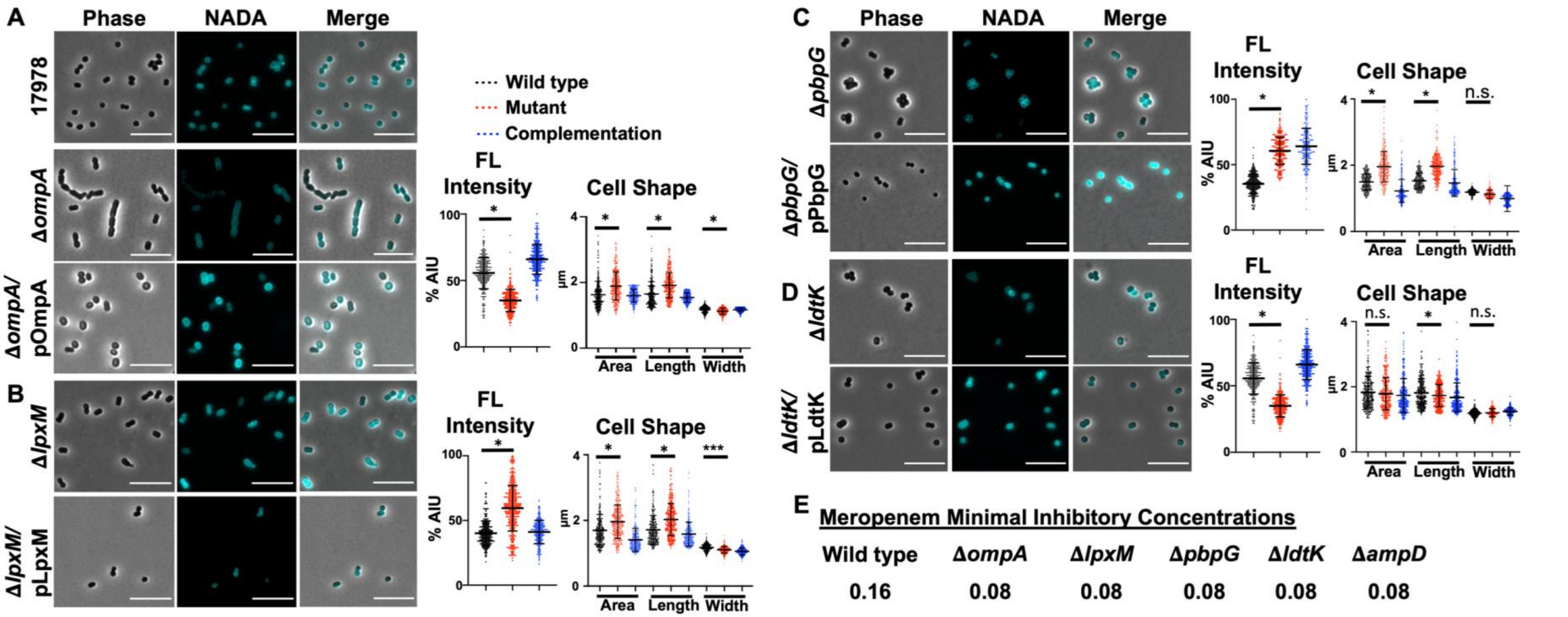

**Figure S3: Morphology of wild type and mutants *A. baumannii* strains.** (A) Phase and fluorescence microscopy of NADA-treated wild type and **Δ***ompA*, (B) **Δ***lpxM*, (C) **Δ***pbpG* and (D) **Δ***ldtK* with each respective complementation strain. Scale bar is 10 μm. Fluorescent (FL) signal intensity quantification in percent arbitrary intensity units (AIU) in wild type, mutant and complementation strain are to the right of mutant images. Cell shape quantifications including, area (A), length (L) and width (W) ($n$= 300) are also included. Significance was determined using an unpaired t-test ($P$ <0.05). An asterisk indicates significant differences between wild type and mutant ($P$ <0.05); n.s., not significant. Error bars indicate standard deviation. (E) Minimal inhibitory concentrations of wild type and mutant strains.

**Figure S4**

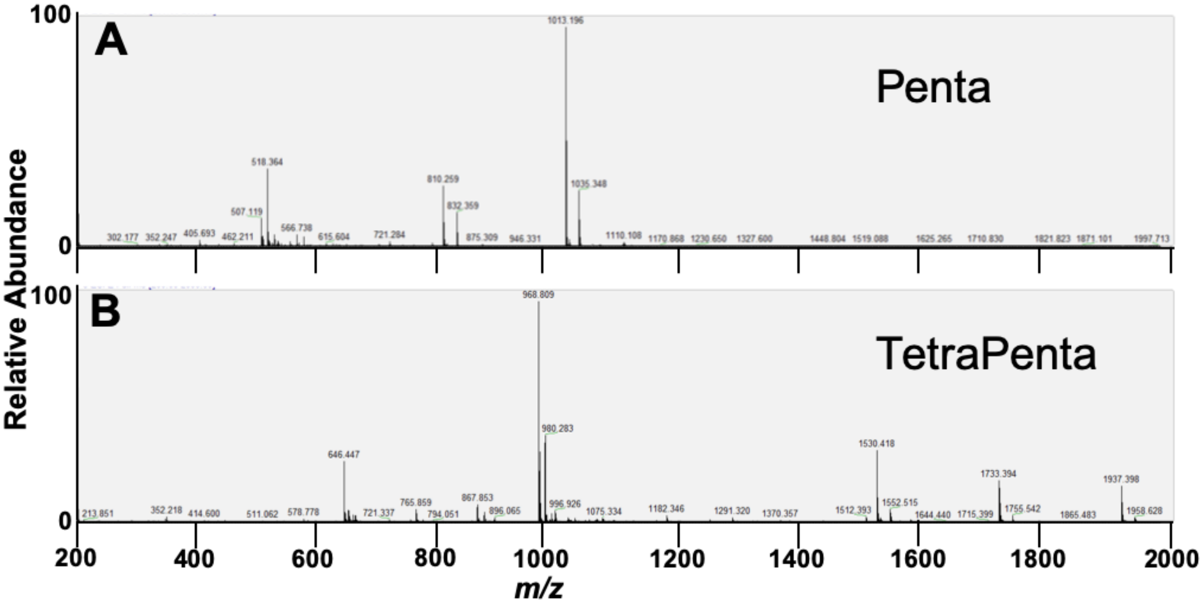

**Figure S4: Unidentified peaks in logarithmic growth phase of Δ*pbpG*.** Peaks at (A) 42 min and (B) 72 minutes were analyzed by mass spectrometry (MS). The peak in A was consistent with disaccharide pentapeptide (Penta, neutral mass: 1012.19 amu; theoretical: 1012.45 amu) and the peak in B was consistent with bis-disaccharide tetrapentapeptide (TetraPenta, neutral mass: 1935.60 amu; theoretical: 1935.84 amu).

**Figure S5**

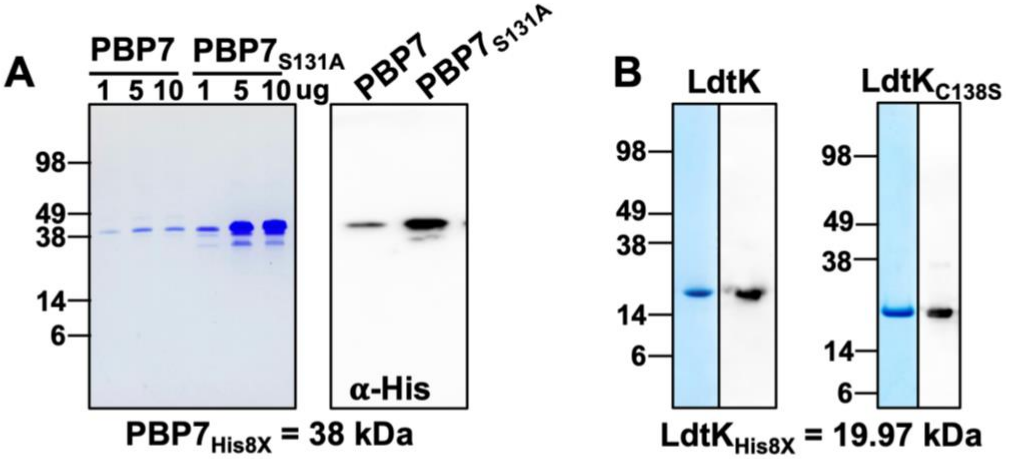

**Figure S5: (A) Purification of recombinant PBP7 and LdtK.** (A) $PBP7_{His8X}$ Left: Coomassie stained SDS-PAGE gel with 1, 2 or 10 µg of recombinant $PBP7_{His8x}$ or $PBP7_{S131A\ His8X}$ loaded. Right: Western blot of 1 µg $PBP7_{His8x}$ or $PBP7_{S131A\ His8X}$ using an **α**-his antibody. (B) $LdtK_{His8X}$ Coomassie stained SDS-PAGE gel with 10 µg of recombinant $LdtK_{His8x}$ or $LdtK_{S131A\ His8X}$ loaded (left) and western blot with 1 µg of each recombinant protein using an **α**-pentahis antibody (right).

### Supplementary Tables

#### Table S1: Muropeptide composition of wild type and Δ*pbpG A. baumannii* strain ATCC 17978.

| Peak No. | Muropeptide | Relative % of each muropeptide[a] | | | |
|---|---|---|---|---|---|
| | | WT Logarithmic | WT Stationary | Δ*pbpG* Logarithmic | Δ*pbpG* Stationary |
| **1** | **Tri** | 3.0 ± 0.4 | 3.2 ± 0.0 | 1.5 ± 0.2 | 2.0 ± 0.1 |
| **2** | **Tri-D-Asn** | 0.9 ± 0.1 | 0.1 ± 0.2 | 0.1 ± 0.3 | 0.1 ± 0.2 |
| **3** | **Tri-D-Lys** | 2.2 ± 0.7 | 4.8 ± 0.0 | 1.0 ± 0.0 | 1.1 ± 0.2 |
| **4** | **TetraGly4** | 0.0 ± 0.0 | 0.7 ± 0.1 | 0.0 ± 0.0 | 0.0 ± 0.0 |
| **5** | **Tetra-D-Lys** | 0.0 ± 0.0 | 2.6 ± 0.0 | 0.0 ± 0.0 | 0.0 ± 0.0 |
| **6** | **Tetra** | 17.6 ± 0.5 | 17.5 ± 0.4 | 16.1 ± 0.2 | 14.6 ± 0.3 |
| **7** | **Tetra-D-Arg** | 0.0 ± 0.0 | 0.4 ± 0.0 | 0.0 ± 0.0 | 0.1 ± 0.2 |
| **7B** | **Penta** | 0.0 ± 0.0 | 0.5 ± 0.0 | 1.3 ± 0.0 | 0.4 ± 0.0 |
| **8** | **TetraTriDapGly4** | 0.3 ± 0.0 | 0.8 ± 0.0 | 0.0 ± 0.0 | 0.2 ± 0.0 |
| **9** | **TriTri(Dap)/TriTriDap-D-Lys** | 0.5 ± 0.1 | 0.5 ± 0.0 | 0.1 ± 0.2 | 0.0 ± 0.0 |
| **10** | **TetraTri(Dap)** | 0.0 ± 0.0 | 1.8 ± 0.1 | 0.0 ± 0.0 | 0.4 ± 0.0 |
| **11** | **TetraTri** | 4.3 ± 0.2 | 3.4 ± 0.1 | 3.1 ± 0.0 | 1.9 ± 0.0 |
| **12** | **TetraTri-D-lys** | 0.6 ± 0.0 | 3.7 ± 0.0 | 0.5 ± 0.0 | 1.0 ± 0.0 |
| **13** | **TetraTri-D-lys** | 0.4 ± 0.0 | 0.7 ± 0.0 | 0.4 ± 0.0 | 0.0 ± 0.0 |
| **14** | **TetraTri-D-Arg** | 0.5 ± 0.1 | 6.0 ± 0.0 | 2.0 ± 0.0 | 1.7 ± 0.0 |
| **15** | **TetraTetra** | 36.2 ± 0.9 | 22.9 ± 0.4 | 39.6 ± 0.6 | 35.7 ± 0.2 |
| **15B** | **TetraPenta** | 0.5 ± 0.1 | 1.0 ± 0.1 | 2.9 ± 0.0 | 1.0 ± 0.1 |
| **16** | **TetraTetraTri or TetraTetraTriDap** | 0.1 ± 0.0 | 0.9 ± 0.0 | 0.9 ± 0.0 | 1.2 ± 0.1 |
| **17** | **TetraTetraTri or TetraTetraTriDap** | 0.3 ± 0.0 | 1.6 ± 0.0 | 0.3 ± 0.2 | 0.4 ± 0.0 |
| **18** | **TriTriDap-D-Met** | 0.6 ± 0.0 | 0.5 ± 0.0 | 0.2 ± 0.4 | 0.6 ± 0.0 |
| **19** | **TetraTetraTetra** | 17.9 ± 0.0 | 11.8 ± 0.3 | 16.9 ± 0.2 | 18.6 ± 0.1 |
| **20** | **TetraTri-D-Met** | 0.1 ± 0.2 | 0.7 ± 0.0 | 0.7 ± 0.0 | 0.2 ± 0.0 |
| **21** | **TetraTri Anh / TetraTetraTetraTri** | 4.4 ± 0.1 | 2.4 ± 0.2 | 3.9 ± 0.0 | 4.7 ± 0.0 |
| **22** | **TetraTetra Anh I** | 1.4 ± 0.0 | 0.8 ± 0.0 | 2.0 ± 0.0 | 2.2 ± 0.1 |
| **23** | **TetraTetra Anh II** | 0.7 ± 0.1 | 0.9 ± 0.0 | 1.0 ± 0.0 | 1.4 ± 0.1 |
| **24** | **TetraTetraTetra Anh** | 2.5 ± 0.1 | 1.6 ± 0.0 | 2.2 ± 0.0 | 3.2 ± 0.0 |
| **Sum of known peaks** | | 95.8 ± 0.2 | 91.7 ±0.1 | 95.8 ± 0.2 | 92.8 ± 0.4 |
| **Monomers (Total)** | | 24.8 ± 0.6 | 32.3 ±0.3 | 20.9 ± 0.0 | 19.7 ± 0.0 |
| **Monomers with modification** | | 3.2 ± 0.6 | 9.3 ±0.2 | 1.1 ± 0.3 | 1.4 ± 0.3 |
| **Monomer tri** | | 3.2 ± 0.5 | 3.5 ±0.0 | 1.6 ± 0.2 | 2.2 ± 0.1 |
| **Monomer tri-D-Asn** | | 0.9 ± 0.1 | 0.1 ±0.3 | 0.1 ± 0.3 | 0.1 ± 0.2 |
| **Monomer tri-D-Lys** | | 2.3 ± 0.7 | 5.2 ±0.0 | 1.0 ± 0.0 | 1.1 ± 0.2 |
| **Monomer tetraGly4** | | 0.0 ± 0.0 | 0.8 ±0.1 | 0.0 ± 0.0 | 0.0 ± 0.0 |
| **Monomer tetra-D-Lys** | | 0.0 ± 0.5 | 2.8 ±0.0 | 0.0 ± 0.0 | 0.0 ± 0.0 |
| **Monomer tetra** | | 18.4 ± 0.5 | 19.0 ±0.4 | 16.8 ± 0.2 | 15.7 ± 0.4 |
| **Monomer tetra-D-Arg** | | 0.0 ± 0.0 | 0.4 ±0.0 | 0.0 ± 0.0 | 0.1 ± 0.2 |
| **Monomer penta** | | 0.0 ± 0.0 | 0.5 ±0.0 | 1.4 ± 0.0 | 0.4 ± 0.0 |
| **Dimers (Total)** | | 52.5 ± 0.4 | 50.3 ±0.2 | 58.8 ± 0.2 | 55.1 ± 0.2 |
| **Dimers with modification** | | 2.9 ± 0.5 | 14.1 ±0.1 | 4.1 ± 0.7 | 4.0 ± 0.1 |
| **Dimer chain ends (anhydroMur*N*Ac)** | | 6.8 ± 0.0 | 4.4 ±0.2 | 7.2 ± 0.0 | 9.0 ± 0.1 |
| **Trimers (Total)** | | 22.6 ± 0.2 | 17.4 ±0.5 | 21.2 ± 0.2 | 25.2 ± 0.1 |
| **Trimer chain ends (anhydroMur*N*Ac)** | | 2.6 ± 0.1 | 1.7 ±0.0 | 2.3 ± 0.1 | 3.5 ± 0.0 |
| **Tripeptides (Total)** | | 13.0 ± 1.2 | 20.8 ±0.4 | 9.0 ± 0.7 | 10.1 ± 0.2 |
| **Tripeptides with modifications** | | 5.2 ± 0.9 | 13.0 ±0.3 | 3.4 ± 1.0 | 3.6 ± 0.1 |
| **Tetrapeptides (Total)** | | 86.3 ± 1.3 | 77.5 ±0.4 | 89.0 ± 0.9 | 88.9 ± 0.1 |
| **Tetrapeptides with modifications** | | 0.9 ± 0.2 | 10.5 ±0.1 | 1.9 ± 0.0 | 1.8 ± 0.3 |
| **Pentapeptides** | | 0.2 ± 0.1 | 1.0 ±0.0 | 2.9 ± 0.0 | 1.0 ± 0.1 |
| **3-3 Crosslinks** | | 1.1 ± 0.1 | 2.9 ±0.1 | 0.6 ± 0.3 | 1.2 ± 0.1 |
| **Chain ends (anhydroMur*N*Ac)** | | 4.3 ± 0.0 | 2.8 ±0.1 | 4.3 ± 0.0 | 5.7 ± 0.1 |
| **Degree of crosslinkage** | | 41.4 ± 0.3 | 36.7 ±0.2 | 43.6 ± 0.0 | 44.3 ± 0.0 |
| **% peptides in cross-links** | | 75.2 ± 0.6 | 67.7 ±0.3 | 79.1 ± 0.0 | 80.3 ± 0.1 |

**[a]Values are mean ± variation of two biological repeats.**

**Table S2**

**Table S2:** Strains and plasmids used in this study

| Strain/Plasmid | Description | Reference/Source |
|---|---|---|
| **Strains** | | |
| *E. coli* C2987 | chemically competent wild type, K-12 | New England Biolabs |
| *E. coli* C2527 | chemically competent BL-21 | New England Biolabs |
| *A. baumannii* ATCC 17978 | wild type | ATCC (1) |
| *A. baumannii* ATCC 19606 | wild type | ATCC (2) |
| *A. baumannii* AYE | wild type | ATCC (3) |
| *A. baumannii* AR Bank # 273 | Clinical isolate | (4) |
| *A. baumannii* AR Bank # 280 | Clinical isolate | (4) |
| *A. baumannii* AR Bank # 300 | Clinical isolate | (4) |
| *A. baumannii* ATCC 17978 | Δ*ompA* | This Study |
| *A. baumannii* ATCC 17978 | Δ*ompA* / pOmpA | This Study |
| *A. baumannii* ATCC 17978 | Δ*lpxM* | (5) |
| *A. baumannii* ATCC 17978 | Δ*lpxM* / pLpxM | (5) |
| *A. baumannii* ATCC 17978 | Δ*pbpG* | This Study |
| *A. baumannii* ATCC 17978 | Δ*pbpG* / pPBP7 | This Study |
| *A. baumannii* ATCC 17978 | Δ*ldtK* | (6) |
| *A. baumannii* ATCC 17978 | Δ*ldtK* / pLdtK | (6) |
| *A. baumannii* ATCC 17978 | Δ*ampD* | This Study |
| *A. baumannii* ATCC 17978 | Δ*ampD* / pAmpD | This Study |
| **Plasmids** | | |
| pMMB67EH | Amp$^R$ | (7) |
| pABBR | Amp$^R$ | (8) |
| pABBRKn | pABBR_MCS with the *Kan*$^R$ gene from pKD4 inserted into the PvuI site, Kn$^R$ | (9) |
| pMMB67EHKn | pMMB67EH with the *Kan*$^R$ gene from pKD4 inserted into the PvuI site, Kn$^R$ | (6) |
| pJNW684 | Tn vector, Amp$^R$, Kan$^R$ | (10) |

| | | |
|---|---|---|
| pAT03 | pMMB67EH with FLP recombinase, Amp$^R$ | (8) |
| pAT04 | pMMB67EH with REC$_{Ab}$ system, Tet$^R$ | (8) |
| pKD4 | Kan$^R$ | (11) |
| pT7-7 | Amp$^R$ | (12) |
| pT7-7Kn | pT7-7 with the *Kan$^R$* gene from pKD4 inserted into the PvuI site, Kn$^R$ | This study |
| pUC19 | Amp$^R$ | (13) |
| pLdtK | pMMB67EHKn with the *ltdK* (A1S_2806) gene and IPTG inducible promoter inserted into the KpnI and SalI sites, Kn$^R$ | (6) |
| pUC19::LdtK$_{C138S}$ | pUC19 with *ltdK*$_{C138S}$ (A1S_2806) gene cloned into the BamHI sites, Amp$^R$ | (6) |
| pLdtK$_{C138S}$ | pMMB67EHKn with the *ltdK*$_{C138S}$ (A1S_2806) gene and IPTG inducible promoter inserted into the KpnI and SalI sites, Kn$^R$ | (6) |
| pLdtK-His$_8$-tag | pT7-7 with *ltdK*$_{C138S}$ (A1S_2806) cloned into the NdeI and BamHI sites, Kn$^R$ | This study |
| pLdtK$_{C138S}$-His$_8$-tag | pT7-7 with *ltdK*$_{C138S}$ (A1S_2806) cloned into the NdeI and BamHI sites, Kn$^R$ | This study |
| pPBP7 | pMMB67EHKn with the *pbpG* (A1S_0237) gene and native promoter inserted into the XhoI and KpnI sites, Kn$^R$ | This study |
| pUC19::PBP7$_{S131A}$ | pUC19 with *pbpG*$_{S131A}$ (A1S_0237) gene cloned into the BamHI sites, Amp$^R$ | This study |
| pPBP7-His$_8$-tag | pT7-7 with *pbpG* (A1S_0237) cloned into the NdeI and BamHI sites, Kn$^R$ | This study |
| pPBP7$_{S131A}$-His$_8$-tag | pT7-7 with *pbpG*$_{S131A}$ (A1S_0237) cloned into the NdeI and BamHI sites, Kn$^R$ | This study |
| pOmpA | pMMB67EH with *ompA* (A1S_2840) cloned into the KpnI and SalI sites, Amp$^R$ | This study |
| pLpxM | pMMB67EH with *lpxM* (A1S_2609) cloned into the KpnI and SalI sites, Amp$^R$ | (5) |

**Table S3**

### Table S3: Oligonucleotides used in this study

| Oligo Name | Sequence (5' to 3') |
|---|---|
| **Deletion Primers** | |
| 17978 *ompA* Kan-FRT 5' | TCAAGTGTTTGTATGATTCAAATGTGAATAGCTTAAAAATAATACTGGGGTAAAAAAATATCTCAGGGGCCAATAAATTTAGGCTGAGCTTGAACAACAATTGTTATCTCTGGAGGATATCCATGagcgattgtgtaggctggagctgcttcg |
| 17978 *ompA* Kan-FRT 3' | ATTTTAAGTAATGATTGGAAGAGATTATGAATCAGGAGATTTACAAATGACCAAATATTTTAAAAATCGCCATAAAAAAAGCGACTCTAACGAGTCGCTTTTTTACTGTTCAAGAACTCAAATTAatatcctccttagttcctattccg |
| 17978 *ompA*-kn verify 5' | ACTGGTAAAATCACGGCAAGCG |
| 17978 *ompA*-kn verify 3' | CCTCGTGCTCAACATCGTAGAAATAG |
| 17978 *ampD* Kan-FRT 5' | ATTAACTGTTTTAAACTAAAACAATTGGCAAGCAATTTATTGGCATCTCAATGAAGCAAATCACACCGTATGAAGTTATAGATGGACAATTAAAAGGGGCGAGACAAGTACCTTCTCCAAATTTTAGCGATTGTGTAGGCTGGAGCTGCTTCG |
| 17978 *ampD* Kan-FRT 3' | TGCCATTTAAAATAAGGTCCCGGATCGGTTTTTCGGCCCGGTGCAATGTCTGAATGCCCTGCAATATGGTTTTTAATTTCAGGATAGGCCTGACGAATAGCAGTAACAACGTCAGTTAGCACTTCATATCCTCCTTAGTTCCTATTCCG |
| 17978 *ampD*-kan verify 5 | GACTGAGGTATGCTCATACGG |
| 17978 *ampD*-kan verify 3 | CGAAATCTTTACCCGCACCAAA |
| 17978 *pbpG*-Kn FRT F | AAAGCTTTATACCTTATATCTCAAATGTAAGGCATAATGATAGTAAGCGCAAATGTGTGTCACCCTGAGTCGAGTATTGTGCCGTGAAAAATTCTAAAAAGTCTTTAATGCATGTGCTAAGCATGATATCCTCCTTAGTTCCTATTCCG |
| 17978 *pbpG* -Kn FRT R | TATCTAATATGTAAAATCTGAGTTTTTATAAAAAGCGGCTGTTTAATACAGCCGTTTTTTTATGCTTTTTAAATGGCATAAAAAAACGCTTCTTAAAAGAAGCGTTTTTAAAAATAATTAAATTAAGCGATTGTGTAGGCTGGAGCTGCTTCG |
| 17978*pbpG* confirm Up | GCTTGCAATGGAATGACAAAATTAGCAATC |
| 17978*pbpG* confirm Dwn | GATACAGCAATTAAACAATGTGCTGATGCAG |
| **Complementation Primers** | |
| 17978 o*mpA* CDS KpnI | CGCGGTACCATGAAATTGAGTCGTATTGCACTTGCTAC |
| 17978 *ompA* CDS SalI | CGCGTCGACTTATTGAGCTGCTGCAGGAGCTGCC |
| 17978 *ampD* CDS BamHI | CGCGGATCCATGAAGCAAATCACACCGTATGAAGTTATAGAT |
| 17978 *ampD* CDS SalI | CGCGTCGACTCAAGTTTTTTTCTGTGCTAACAACTGCCTAAA |
| pABBR-*pbpG*-F XhoI | AAATTACTCGAGCTATTCTTCTATAGTGAGCGAATAGTTG |
| pABBR-*pbpG*-R KpnI | GTTGTCGGTACCTGCAACAATGGACCAAGTAAAAGATTCG |
| **Mutagenesis Primers** | |

| | |
|---|---|
| *pbpG*$_{S131A}$ Fwd | CAGGTGAGGTACTTTATAGTAAAAATACCAACGCATCAGTGCCGATCGCTGCAATTACCAAATTGATGACGGCAGTTGTAACGGCAGATGCCCGTTTAAACAT |
| *pbpG*$_{S131A}$ Rev | ATGTTTAAACGGGCATCTGCCGTTACAACTGCCGTCATCAATTTGGTAATTGCAGCGATCGGCACTGATGCGTTGGTATTTTTACTATAAAGTACCTCACCTG |
| pUC19-*pbpG* BamHI 5' | CGCGGATCCATGTCTATTTTGCTTAGT |
| pUC19-*pbpG* BamHI 3' | CGCGGATCCTTAAATACGTTTTGGCAAAT |
| **Overexpression Primers** | |
| pT7-*pbpG* NdeI | CGCCATATGTCTATTTTGCTTAGT |
| pT7-*pbpG* BamHI 8X-his | CGCGGATCCTTAATGGTGATGGTGATGGTGATGGTGAATACGTTTTGGCAAAT |
| **Sequencing Primers** | |
| pMMB67EH seq fwd | CGGTTCTGGCAAATATTCTGAAA |
| pMMB67EH Seq3 | CTGCGTTCTGATTTAATCTGTAT |
| pABBR confirm 1 | GGGCTGACCGCTTCCT |
| pABBR confirm 2 | CGCTAGCAGCACGCCATAG |
